# Evidence of hard-selective sweeps suggest independent adaptation to insecticides in Colorado potato beetle (Coleoptera: *Chrysomelidae*) populations

**DOI:** 10.1101/2022.08.15.504016

**Authors:** Zachary P. Cohen, Yolanda H. Chen, Russell Groves, Sean D. Schoville

## Abstract

Pesticide resistance provides one of the best examples of rapid evolution to environmental change. The Colorado potato beetle has a long and noteworthy history as a super-pest due to its ability to repeatedly develop resistance to novel insecticides and rapidly expand its geographic and host plant range. Here, we investigate regional differences in demography, recombination and selection using whole genome resequencing data from two highly resistant CPB populations in the United States (Hancock, Wisconsin and Long Island, New York). Demographic reconstruction corroborates historical records for a single pest origin during the colonization of the Midwestern and Eastern United States in the mid-to late-19th century and suggests that the effective population size might be higher in Long Island, NY than Hancock, WI despite contemporary potato acreage of Wisconsin being far greater. Population-based recombination maps show similar background recombination rates between these populations, as well as overlapping regions of low recombination that intersect with important metabolic detoxification genes. In both populations, we find compelling evidence for hard selective sweeps linked to insecticide resistance with multiple sweeps involving genes associated with xenobiotic metabolism, stress response, and defensive chemistry. Notably, only two candidate insecticide resistance genes are shared among both populations, but both appear to be independent hard selective sweep events. This suggests that repeated, rapid, and independent evolution of genes may underlie CPB’s pest status among geographically distinct populations.

## Introduction

The evolution of insecticide resistance is considered one of the most pressing issues facing insect pest management and sustainable agricultural practice. Despite considerable research effort over the last 40+ years (Georghiou and Taylor 1977; Roush 1989), evolutionary predictions from insect resistance management models have continued to underperform. In many cases, assumptions underlying these models (*i*.*e*., a single recessive resistance allele, fitness tradeoffs for resistance mutations, and a constant selection intensity from insecticides) might not apply to real-world evolution of insecticide resistance given observations of large population sizes, high standing genetic diversity and accumulation of resistance alleles over time (Hoy 1998; Groeters and Tabashnik 2000; Brévault et al., 2013; Bass et al., 2015). Consequently, pesticide resistance in herbivorous insects is still a major concern for the global food supply, as these pests cause an estimated 18-20% damage (∼$470 billion annually) to crops (Sharma et al. 2017). Research and development for new insecticides is becoming more difficult (Gould et al. 2018), and costs have ballooned by 84.8% since 2008 (McDougall 2011). However, knowledge of insect pest population genetics has been slow to develop, particularly in terms of understanding the genomic mechanisms and population genetic parameters that underlie resistance evolution (Pélissié et al. 2018).

Insecticide resistance phenotypes can be achieved through evolutionary changes in multiple pathways, such as: target-site insensitivity (knockdown resistance), reduced penetration (cuticular change), xenobiotic detoxification and transport (metabolic processing by ABC transporters, cytochrome P450s, glutathione S-transferases, and carboxylesterases), as well as behavioral avoidance of toxic compounds (Scott et al. 1998; Mamidala et al. 2011; ffrench-Constant 2013). As a case in point, the Colorado potato beetle (CPB), *Leptinotarsa decemlineata*, shows evidence of at least two types of physiological resistance: target-site insensitivity and metabolic detoxification (Hawthorne 2003, Alyokhin et al. 2008). The evolution of target-site insensitivity could be attributed to a single locus (Rinkevitch et al., 2012), but metabolic detoxification seems to involve multiple molecular pathways (Clements et al., 2016; Zhu et al. 2016) and draws from different genes in those pathways across geographically dispersed resistant populations (Dively et al., 2020; Pélissié et al., 2022).

CPB is among the most notorious pests, with documented resistance to 56 insecticides (Whalon et al. 2016) spanning all modes of action (IRAC 2016), representing nearly 150 independent reports, and thus is an important model of agricultural pest genomics (Schoville et al., 2018). From early insecticides such as Paris Green and DDT, to modern insecticides such as pyrethroids and neonicotinoids, CPB populations have accumulated resistance over impressively short periods of time, even within one year (∼2-3 generations) of the introduction of novel chemicals (Forgash 1985, Ioannidis et al. 1991, Alyokhin et al. 2008; Brevik et al., 2018). In a species-level comparative genomic analysis, CPB shows a higher rate of positive selection on putative insecticide resistant loci compared to other species in the genus *Leptinotarsa* (Cohen *et al*. 2020). Recent population genomic sequencing studies of CPB have found evidence for selection acting on many insecticide resistance pathways, supporting a prominent role of polygenic adaptation to insecticides over time (Crossley et al, 2017; Pélissié et al., 2022).

Positive selection appears to act repeatedly on the same molecular pathways (and sometimes the same genes) across geographically isolated populations (Pélissié et al., 2022), but the signature of selection appears to be soft sweeps of polygenic traits on multiple haplotype backgrounds. Hence, these recent studies of CPB provide limited support for the classical insecticide resistance model that involves a single *de novo* mutation. Instead, results suggest that repeated pesticide management failures arise from high standing genetic diversity and the polygenic genomic architecture of resistance traits, which can reduce the waiting time and likelihood of adaptation (Yeaman 2022). If, in fact, field-evolved resistance to insecticides occurs independently among geographically isolated CPB populations and arises from distinct genetic variation, employing general control strategies based on mutations in key target site genes will not lead to effective and sustainable control strategies at the population level.

These previous research efforts were based primarily on genome scans that leverage population differences in allele frequency to identify signatures of selection (Chen et al., 2010), but had limited power to detect hard selective sweeps due to the quality of genomic datasets, *e*.*g*. contiguity, N50, number of scaffolds, etc. Recently, Cohen et al. (2021) generated a near-chromosomal assembly of CPB, enabling improved analysis of linked variation along the genome. These advances motivated this investigation of selective sweeps and provided better inference of the demographic history and genome-wide recombination patterns that contribute to genetic diversity and may confound signals of selection (O’Reilly et al., 2008).

CPB’s native range extends from southern Mexico throughout the plains and high desert regions of the United States (Jacques 1988). Considered to be polyphagous, the Colorado potato beetle can survive on numerous Solanaceous plants, but its primary natal host is buffalo burr, *Solanum rostratum*. Potato beetle larvae and adults both defoliate plants and are therefore equally targeted by insecticides. The larval stage consumes nearly four times more leaf tissue than the adult stage (Pelletier et al., 2011; Ferro et al. 1985). Despite this expansive native distribution, it wasn’t until the mid-19^th^ century that CPB expanded onto cultivated potato, *S. tuberosum*, whereupon it quickly spread eastward. The outbreak onto potato also introduced CPB to other cultivated Solanaceae, such as tomato, eggplant, and tobacco, where it is a minor pest (Jacques 1988). During this rapid expansion, CPB quickly overwhelmed crops and became the target of early chemical control efforts and has remained a key species driving insecticide development and management practices (Alyokhin et al., 2015). We therefore focused our analyses on populations from Hancock, WI and Long Island, NY, two well-studied, highly resistant populations that are geographically distinct and represent different stages in the geographical expansion of pest populations in the United States. Wisconsin potato feeding CPB are some of the first pest populations that migrated (1867) into the state following a major outbreak that started in Omaha, Nebraska (1859) and had ravaged neighboring Iowa by 1862 (Tower 1906). These populations were observed to migrate eastward along a potato growing corridor and ultimately reached the Atlantic seaboard, and Long Island, NY, by 1872. They then invaded continental Europe and eventually spread to Asia (Tower 1906; Grapputo et al., 2005). The CPB populations of Long Island represent the eastern expansion limit in the U.S. from the mid-19^th^ century outbreak and are notoriously pestiferous. Historic eye-witness accounts describe how this population was a major nuisance due to its incredible size. Beetles would spill into Long Island Sound, floating in large mats that would swarm onto the hulls of anchored ships covering the decks and even causing noxious odors for beachgoers when they were washed ashore dead (Tower 1906). The outbreak stopped a train at New York Central Railroad due to the potential hazard of their carcasses slicking the tracks. More recently, the Long Island population demonstrates cross-resistance to multiple insecticides and the most rapid rate of resistance evolution observed in any CPB population (Alyokhin et al., 2015; Dively et al., 2020). Therefore, estimating genomic diversity and reconstructing the demographic history of these iteratively established pest-populations is relevant to determine how colonization history and genomic diversity patterns influence contemporary resistance.

Levels of genome-wide genetic variation are influenced by mutation, recombination, and selection, in direct relationship to effective population size (Hartl and Clark 1997). Prior work suggests that CPB has relatively high genome-wide standing genetic variation (Schoville et al., 2018; Cohen et al., 2020; Pélissié et al., 2022), yet it remains unclear how population history, recombination, and selection interact to shape this variation. Here, we examine the influence of demography and recombination on selection between two highly resistant geographically distinct pest CPB populations in the United States. We do this by estimating each populations’ demographic history, generating population-specific recombination maps, and testing for evidence of hard selective sweeps. We then compare the recombination hotspots to hard-sweep regions within and between populations to assess the relative importance of novel mutations driving the evolution of insecticide resistance.

## Methods

### Study Sites and Sampling

In order to determine how adaptation to insecticides has evolved in the Colorado potato beetle, we compared selection and recombination for two CPB populations known to be highly resistant to the neonicotinoid insecticide imidacloprid. Field-collected beetles representing distinct geographical regions from: Hancock, Wisconsin (HAN: N 44.119753, W -89.535683) and Long Island, New York (LI: N 40.905657, W -72.752664) were collected at single sites, in the same growing season of 2015 (Supplemental Table S1). Populations near Hancock Agricultural Research Station have been studied primarily due to their relatively high resistance to neonicotinoid (Imidacloprid®) insecticides (Huseth et al., 2014; Clements et al., 2016; Crossley et al., 2017), but they are also known to be resistant to several insecticidal chemistries (Dively et al., 2020).

In this study, we generated novel whole genome resequencing data comprising 43 CPB adults (NCBI PRJNA753140), while combining it with previously published data from 10 adults (Accessions: SRR10388315 - SRR10388319, SRR10388359 - SRR10388363), for a total of 53 beetles (N_HAN_ = 28, N_LI_ = 25). For all samples, high molecular weight genomic DNA was isolated from thoracic muscle tissue using DNeasy Blood & Tissue kits (Qiagen).

### Whole Genome Sequencing and Variant Calling

Whole genome sequence reads were generated at the University of Wisconsin Biotechnology Center, using either the Illumina HiSeq 2500 to generate 2 × 125 bp reads (previously published samples) or the Illumina NovaSeq 6000 platform to generate 2 × 150 bp reads (newly incorporated samples). Sequencing libraries were prepared using TruSeq DNA kits (Illumina). Sequencing effort was designed to yield >5x average coverage for each sample, a quantity sufficient to identify SNPs with reasonable accuracy (Li et al. 2009). Samples were demultiplexed, adapters removed, and reads trimmed to remove low quality base calls using BBMap v38.7 (https://sourceforge.net/projects/bbmap/). Paired-end read data was mapped to an updated reference genome from a non-pest individual sampled from Holly, Colorado (NCBI PRJNA750038) using BWA-mem v0.7.17-r1188 (Li et al., 2013). Identification of polymorphic sites was carried out using ANGSD (Korneliussen et al., 2014), while applying filters for quality (at least 3x coverage per individual, with minimum and maximum total coverage across all individuals of 75x and 250x, respectively), base quality (-minMapQ 30), minor allele frequency (> 0.05), and genotype probability (p-value less than 1e-6). Given the different sequencing platforms used to generate our read data we tested for biases in allele frequency, which could influence our variant detection (De-Kaye et al 2020), by sub-sampling the population vcfs by platform, randomizing samples, and testing for significant differences using a one-way ANOVA in Excel v16.6.

### Demographic Reconstruction and Split Time Analysis

In order to reconstruct demographic history and estimate divergence time between these populations, we used two demographic reconstruction approaches based on the site frequency spectrum (SFS). Stairway plot 2 (Liu and Fu 2020) was used to estimate recent demographic history from each population using the folded site frequency spectrum generated in ANGSD. The dataset was restricted to neutral, intergenic sequences, under a setting where singletons were removed due to potential sequencing errors in the low coverage data (Supplemental Methods).

We assumed two generations per year from observations of pest CPB voltinism at these latitudes (Jolivet et al., 1988) and, due to the lack of a specific CPB or related beetle mutation rate, we employed a recently determined mutation rate of 2.1e^-9^ from the non-biting midge, *Chironomus riparius*, which falls within the range of many insect species (Oppold et al., 2017). Coleoptera and Diptera are sister groups that diverged from one another ∼380MYA (Thomas et al., 2020).

The second demographic method employed was based on a joint or 2D site frequency spectrum in dadi v2.1 (Gutenkunst et al., 2009). We used the dadi_pipeline v3.1.6 with model optimization (Portik et al. 2017) to reconstruct the split time for HAN and LI populations. A combined 2-dimensional site frequency spectrum, 2D-SFS, was generated from intergenic regions with a minor allele frequency of 5% in easySFS (https://github.com/isaacovercast/easySFS). We used dadi_pipeline to compare two probable models of demographic history: 1) a divergence model with no migration and constant population size (no_mig), and 2) a divergence model with no migration, but allowing for population size fluctuation in the descendant lineages (no_mig_size). The models were defined based on current knowledge of the beetles’ dispersal ability, as ecological dispersal rates are on average less than ∼500 meters per year (Boiteau et al., 2003) and genetic data suggests populations do not interbreed over large distances (Grapputo et al., 2005; Izzo et al., 2018). These models were subsequently tested with four consecutive rounds of optimization. Multiple replicates and parameter settings were used at each round with the default settings of dadi_pipeline (replicates = 10, 20, 30, 40; maxiter = 3, 5, 10, 15; fold = 3, 2, 2, 1), and parameters were optimized using the Nelder-Mead method (optimize_log_fmin). Replicate results were summarized using the Summarize_Outputs python script from dadi_pipeline and model selection was based on the Akaike information criterion (AIC; Akaike 1973). To confirm that sample size (N=53) did not influence model selection, a corrected AIC (AICc) was calculated for each dadi model using parameters based on Hurvich and Tsai (1989). Ancestral population size, N_a_, and divergence between Wisconsin and New York were calculated using the formula *θ* = *4?*_*a*_*μL*, where theta (*θ*) is provided from dadi output, mu (*μ*) is the midge mutation rate of 2.1e^-9^, and the length (*L*) in base pairs includes all intergenic sequence data (∼840Mb). After solving for N_a,_ this value is multiplied by divergence time (*T*) and divided by the average number of CPB generations per year (2) to get approximate divergence in years. These parameters were averaged among the five replicates per model.

### Estimation and Comparison of Population Recombination Maps

We generated fine-scale recombination maps per scaffold per population in order to compare rates between populations and correlate selectively swept regions with contiguous haplotypes, *i*.*e*., low recombining regions via pyrho v0.1.6 (Spence and Song 2019. This program uses a composite-likelihood approach to infer recombination maps from individual polymorphism data. The genotype data are used to compute a lookup table of two-locus likelihoods of linkage disequilibrium, which are used to bound the hyperparameters of the model. An innovative feature of pyrho is that the lookup table accounts for demographic change by using independent estimates of effective population size over time while computing the two-locus likelihoods.

To determine how population demography might support the single origin hypothesis for pest CPB populations, we used demographic estimates for each population from stairway plot 2 (see above). Scaffold-specific population vcfs were used to estimate recombination in pyrho using a window size of 80 SNPs, with a step value 15 SNPs, for both populations. These parameters were determined from the pyrho_hyperparam function, while other parameters were left in default settings. The output from pyrho is an estimate of the per generation recombination rate per base (r). To convert to population recombination rates, we used the formula ρ = 4N_e_ r for autosomal scaffolds, where N_e_ is the effective population size estimate for each population from dadi. For the X-chromosome scaffold (CPB has an XO system, where males lack a Y chromosome), we assumed no sex bias in the contribution to N_e_ and multiplied the autosomal effective size (4N_e_) by 3/4 (Wright 1984), while we adjusted the recombination rate (r) by ? as recombination on the X chromosome only occurs in females (Lohmueller et al., 2010). The mean values of “r” per population for the entire genome were compared for significant difference using a Student’s t-Test in R v4.1.2 (2021). Extreme high and low recombination rates (>10 fold) were determined across the genome, relative to the average recombination rate of each scaffold. Recombination was also compared between coding and non-coding regions.

### Selective Sweep Identification

Hard selective sweeps, which are named for their quick rise in frequency in a population, homogenous haplotype, and implicit fitness advantage, were identified for each population using the program RAiSD (Alachiotis and Pavlidis 2018), with default parameter settings. Specifically, this program examines several selective sweep signatures across genomic windows, with a step size of 1 SNP, by measuring 1) high and low frequency derived alleles, 2) localized LD on each flank of candidate a sweep locus, and 3) low LD between these flanking regions. RAiSD incorporates these metrics and generates a composite μ statistic that measures sweep intensity to exclusively identify hard sweeps. The analysis was run for each population, and the resulting outlier windows (highest μ values 0.05%) were filtered using a conservative threshold (α = 0.05). To assess the false positive rate for selective sweeps at this threshold, we used neutral simulations generated in the program ms (Hudson 2002). Neutral instantaneous population growth was modeled using parameters estimated for Wisconsin and New York, including estimated effective size (N_NY_∼ 40,000 & N_WI_ ∼15,000), divergence time converted to theta time units, MAF filtering of 5%, recombination rate, and mutation rate. The resulting ms output was then used in RAiSD to determine thresholds for significant SNP outliers at 5%, 0.5%, and 0.05% cutoffs per population (Supplemental Methods).

Significant hard sweep regions were then annotated for overlap with coding regions using the intersect function in bedtools v2.26.0 (Quinlan and Hall 2010). Both SNPs and genes were compared for overlap between the two pest populations. To understand the biological role of candidate genes and if they occurred in shared pathways, we examined the functions of their gene ontology (GO) terms. We conducted a GO enrichment analysis using a Fisher exact test with a hypergeometric distribution, with sampling by non-replacement to assess significance (p < 0.05). Significant GO terms were grouped for similarity using VennPainter v1.2.0 (Lin et al., 2016). Visualization of significant GO terms was done using the web interface Revigo (Supek et al., 2011) and the python program Cirgo (Kuznetsova et al., 2019).

Finally, the distribution of outlier SNPs within shared genic regions was quantified and the size of the window surrounding each outlier region was also measured to support the identification by RAiSD of hard selective sweeps. The window size was taken from the RAiSD results, which reports the start and stop positions encompassing each region flanking a putatively swept SNP. We grouped the results as either hard-swept regions (based on μ) or non-swept regions. We used a non-parametric Wilcoxon signed-rank test (*W*) to calculate differences in the size of hard and non-swept regions.

## Results

### Demographic Reconstruction and Split Time Analysis

After cleaning the reads and curating polymorphisms for quality, the final call set retained 11.8 million and 12.4 million polymorphic sites for Hancock, WI and Long Island, NY populations, respectively of an approximately 870MB genome (**Supplemental Table S1**), with no difference between sequencing platform (pval_WI_ 0.334833941; pval_LI_ 0.133881306; **Supplemental Table S2**). Stairway plot reconstruction shows that both the Long Island, NY and Hancock, WI CPB populations have declined relative to their ancestral population sizes (**Figure 1**). The stairway plot estimated that the founding population for Long Island was about 10 times larger than the Hancock, WI. HAN (∼90%) has declined proportionally more than LI (∼85%), albeit with broad confidence intervals that overlap the range of effective size changes evident in LI. For the LI population, we estimated that the ancestral effective population size plateaued around 150 years ago, whereas the HAN plateaued more recently approximately 90-110 years ago (**Figure 1**). Using a model fitting approach to estimate the population split time, a no migration model with constant population size was preferred over a no migration model with variable size (AIC_no mig_ 86,303.16; AIC_c_ 86,303.56< AIC_no mig size_ 93,710.02; AIC_c_ 93,710.43), with the top five optimized no_mig replicates (**Supplemental Table S3)**. The constant size model was still favored after AICc. Parameter estimation suggested a divergence time of 159 +/-2.65 years ago and effective population size estimates of Ne_LI_= 40,071 +/-1,447 and Ne_HAN_= 14,777 +/-572 individuals. This joint model also suggested growth from a shared founding population of ∼6,700 individuals, with subsequent population growth upon expansion to potato in Wisconsin and New York.

**Figure 1.**
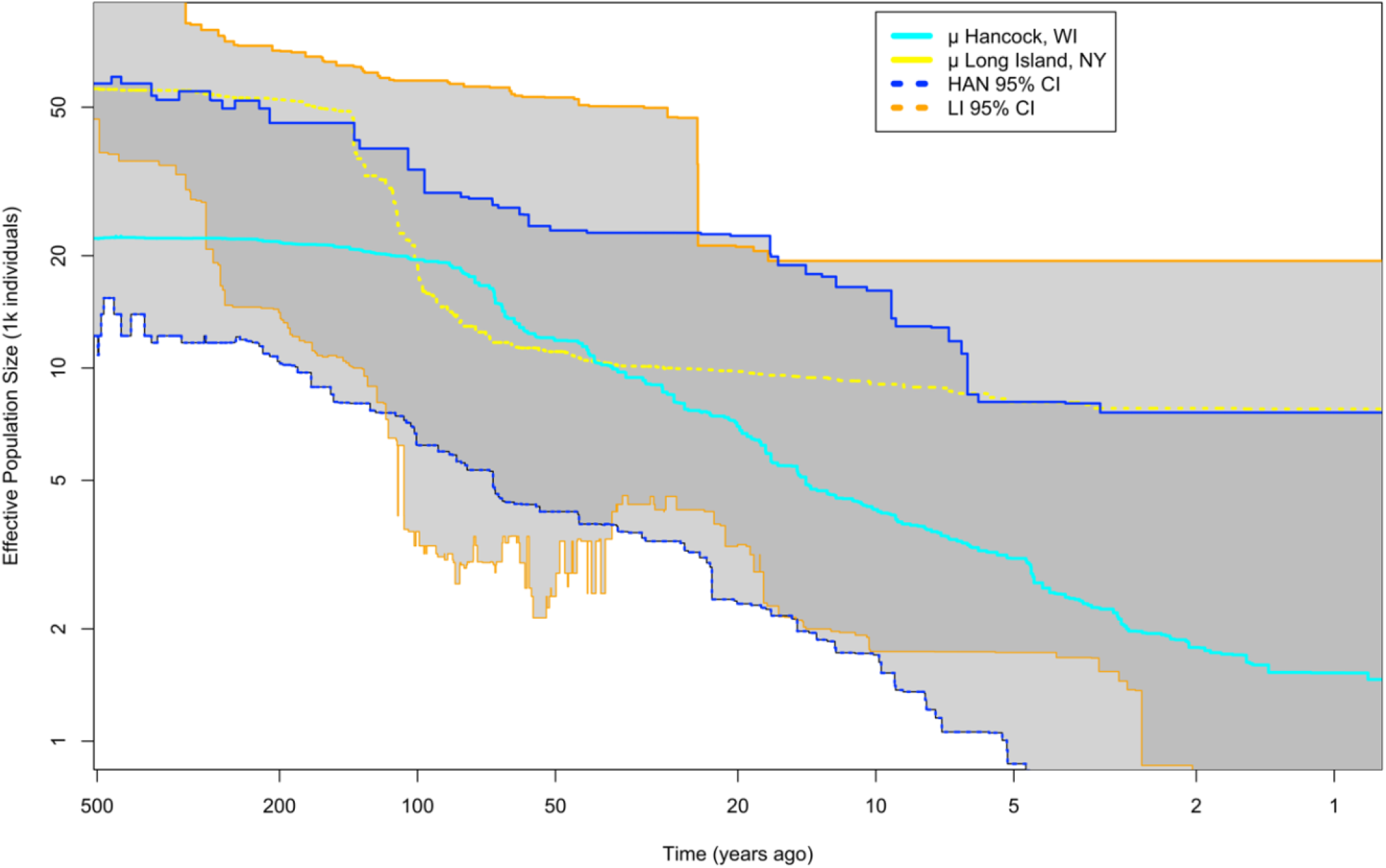
Demographic reconstruction of Colorado potato beetle population size change in Hancock, WI and Long Island, NY. Light shading represents bounds of confidence intervals by population, darker shading is shared confidence intervals (CI) of both populations.

### Estimating Genome-wide Recombination Rates for Each Population

The population-based recombination rate (ρ = 4N_e_ r) for HAN (ρ_HAN_= 0.163) was significantly different at nearly one third of the LI (ρ_LI_ = 0.316) rate. The per-generation per base recombination rate (r) differed only slightly yet was also significantly different (r_HAN_= 2.75e-6 and r_LI_= 1.95e-6; *p-value <2*.*2e-16*). For the X chromosome, the population-based recombination rate (for the X, ρ = 2N_e_r) was lower in HAN (ρ_HAN_=0.086) than LI (ρ_LI_ = 0.195), and much lower than the autosomal rates while the per generation per base rate (r) differed only slightly among populations in comparison to the autosomal rates (r_HAN_= 2.91e-6 and r_LI_= 2.4e-6). Recombination rates reflecting a 10-fold difference above and below the average background scaffold recombination rate were compared among coding and non-coding genomic regions. These rates were slightly higher in genic regions of HAN than non-genic regions (2.78e-6 > 2.69e-6), whereas rates in gene regions for LI were slightly less than non-genic regions (1.9503e-6 < 1.951e-6). The distribution among high and low recombination regions (ten-fold difference from the mean) were compared between populations and examined for gene regions. There were fewer regions of high recombination (LI = 5,295 and HAN = 5,601) than low recombination (LI= 108,549 and HAN = 156,267, see **Figure 2**). A total of 447 genes occurred in shared regions of low recombination, while only three genes occurred in shared regions of high recombination (**Supplemental Table S4**). Notable insecticide resistance genes in shared regions of low recombination include: several cytochrome P450s (CYP9e2, CYP6a23, CYP6a2, CYPb-c1), ABC multidrug resistance-associated proteins, and nAChR subunit α1. One shared gene in a low recombination region (Niemann-Pick type protein, XP_023012615) has evidence of a shared selective sweep (see below) in both LI and HAN. Among the three genes in high-recombining regions, one carboxylesterase gene (XP_023026553, LDEC012644) is potentially related to insecticide resistance.

**Figure 2.**
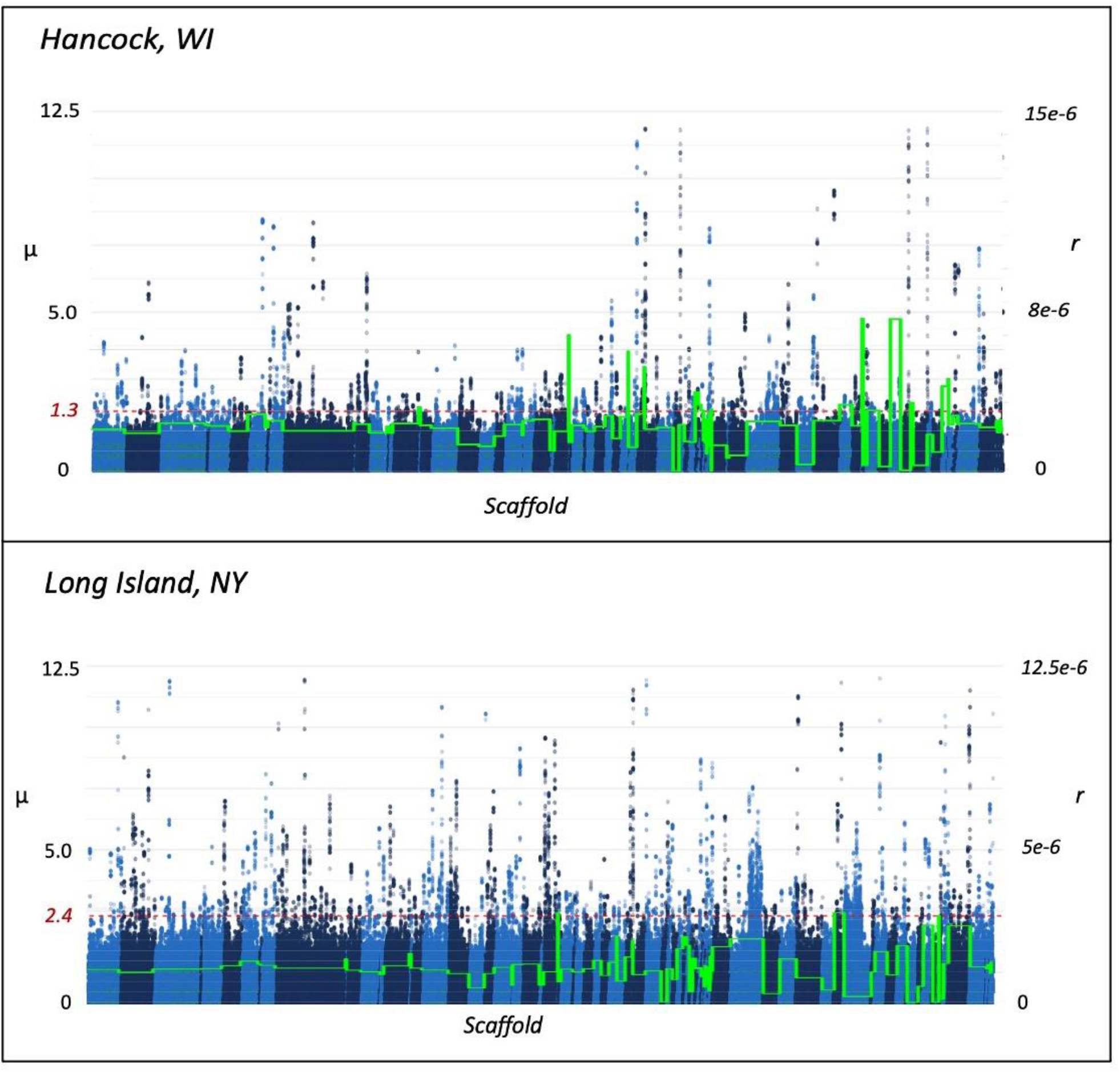
Scaffold averaged recombination of autosomal scaffolds, green trace (r), and selection (μ) with selection significance cutoff (0.05%) for Colorado potato beetle pest populations of Hancock, WI and Long Island, NY.

### Evidence of Shared and Unique Population Hard Selected Sweeps

We examined both populations for evidence of hard selective sweeps using the μ statistic implemented in RAiSD, while assessing thresholds for the false positive rate from neutral simulations (**Figure 3**). A neutral population growth model reconstructed in ms and tested in RAiSD generated μ values slightly larger than observed values, but there was no evidence of very large μ values indicative of hard selective sweeps in the neutral simulations. In contrast, a clear peak of large μ values is evident in the observed data. Defining a conservative 0.05% threshold based on the neutral simulations (μ=2.45e-10), we found approximately 450k SNPs (∼4% of the dataset) significant in Wisconsin and Long Island. Therefore, we opted for the much more conservative 0.05% threshold of observed data for downstream analysis. There were more significant outlier SNPs (6,211) in the Long Island beetle population, with 1,868 of these SNPs located within 89 genes (**Supplemental Table S5**). The HAN population had fewer significant outlier SNPs (5,932), with 1,568 of these SNPs located within 74 genes. Population-specific sweeps include agriculturally relevant genes that could contribute to pest success.

**Figure 3.**
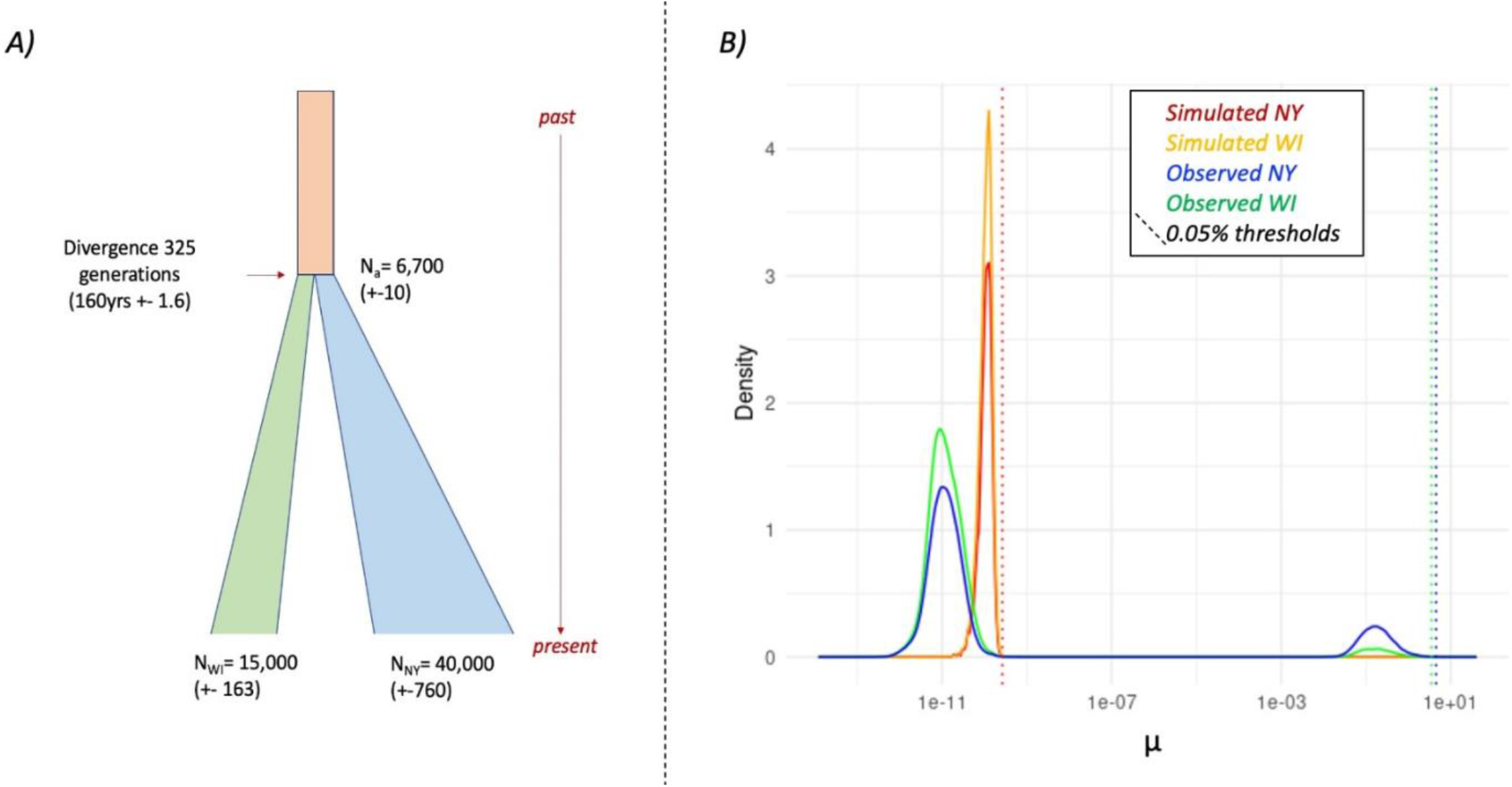
Evidence supporting hard selective sweeps. A) Historical demographic model and parameters used for neutral simulation in ms. B) Simulated and observed selective sweeps from RAiSD (μ statistic, α=0.05) for Colorado potato beetle in Long Island, NY and Hancock, WI. Larger values of μ indicate a hard selective sweep.

The LI and HAN populations shared only nine selectively swept genes, suggesting a minor role for convergent evolution or deeper-time selection on shared ancestry (see **Table 1**), and two of these genes are isoforms of one another, residing within the same genomic region (E3 ubiquitin-protein ligase HECW2 isoform X1 and X2). Among the 89 significant genes in LI, known insecticide resistance loci include: the neuronal acetylcholine receptor subunit alpha-7 (XP_023013985, LDEC018602), cytochrome P450 9e2 (XP_023018113, isoforms X1 and X2), esterase FE4-like (XP_023020747), and the alkaline phosphatase ALP-1 (XP_023023329).

**Table 1.** Shared candidate genes identified as hard selective sweeps in RAiSD in both Hancock, WI and Long Island, NY.

| NCBI gene | Gene annotation | Recombination rate $\rho$ | SNPs | Length of Swept Region (bp) |
| --- | --- | --- | --- | --- |
| XP_023014225 | paired mesoderm homeobox protein 2A-like | 0.227 (WI)<br>0.281 (LI) | 40 (WI)<br>25 (LI) | 24702 (WI)<br>24643 (LI) |
| XP_023018886 | E3 ubiquitin-protein ligase HECW2 isoform X1 | 0.159 (WI)<br>0.244 (LI) | 26 (WI)<br>26 (LI) | 11011 (WI)<br>11959 (LI) |
| XP_023018894 | E3 ubiquitin-protein ligase HECW2 isoform X2 |  |  |  |
| XP_023012615 | Niemann-Pick type protein homolog 1B-like | 0.265 (WI)<br>0.082 (LI) | 9 (WI)<br>2 (LI) | 8551 (WI)<br>9711(LI) |
| XP_023020349 | uncharacterized protein LOC111508933 | 0.176 (WI)<br>0.089 (LI) | 20 (WI)<br>21 (LI) | 20332(WI)<br>16327(LI) |
| XP_023025254 | uncharacterized protein LOC111513291 | 0.181 (WI)<br>0..229 (LI) | 47 (WI)<br>29 (LI) | 42930 (WI)<br>42228 (LI) |
| XP_023025815 | neuroendocrine convertase 2-like | 0.183 (WI)<br>0.405 (LI) | 1 (WI)<br>33 (LI) | 34166 (WI)<br>33273 (LI) |
| XP_023026978 | multidrug resistance-associated protein 1 | 0.194 (WI)<br>0.277 (LI) | 51 (WI)<br>2 (LI) | 61771 (WI)<br>61777 (LI) |
| XP_023027618 | GMP synthase | 0.076 (WI)<br>0.286 (LI) | 33 (WI)<br>23 (LI) | 42468 (WI)<br>42740 (LI) |

Among the 74 significant genes in HAN, known insecticide resistance loci include: the segmentation protein cap-n-collar (XP_023016920), an ATP-binding transporters (ABCD3: XP_023011673 and LDEC007110, ABCC: XP_023026978), salicyl alcohol oxidase (LDEC016943), and an esterase (LDEC022531). Among the nine genes within hard swept regions shared by both populations, there are two candidate insecticide resistance genes: an ABC transporter (ABCC: XP_023026978, LDEC019090) and a carboxylesterase (XP_023025254, LDEC022531).

Gene ontology enrichment resulted in 281 significant terms unique to LI and 169 significant terms unique to HAN, with 27 shared significant terms. While shared enriched GO terms only included a few relevant to insecticide resistance (include calcium ion binding GO:0005509 and transmembrane transport GO:0055085), terms within populations included: HAN: ATPase-coupled transmembrane transporter activity GO:0042626, larval feeding behavior GO:0030536, cellular oxidant detoxification GO:0098869; and LI: ATP-dependent peptidase activity GO:0004176, chloride channel activity GO:0005254, oxidation-dependent protein catabolic process GO:0070407, and voltage-gated cation channel activity GO:0022843 (**Supplemental Figures S1-S7, Supplemental Table S6**).

The distribution of nucleotide variation and length of sweep regions provides information about the pattern of selective sweeps in the genome. SNP abundance in each sweep region and the size of the region was quantified, with HAN having a higher average number of SNPs (23.4/swept locus) compared to LI (19.6/swept locus). The top 0.05% swept regions were significantly larger than non-swept regions for both HAN (HAN swept_u_ = 37,512 >> non-swept_μ_= 3,524; *W =* 1.8e9, *p-value <* 2.2e-16) and LI (LI swept_μ_= 51,138 >> non-swept_μ_ = 3,361; *W* = 2.2e7, *p-value* < 2.2e-16) (**Supplemental Figures S8**). However, the size of the shared sweep regions (that possess a shared significant outlier SNP within 1bp for both HAN and LI) was not significantly larger than the average sweep regions (*W = 84,475, p-value = 0*.*366*).

## Discussion

There is significant interest in understanding how demography and neutral genetic processes influence population diversity and adaptation (Hawkins et al., 2019; Kreiner et al. 2018; Pélissié et al. 2018). An archetype of this phenomenon is rapidly evolving insect pest populations, which can serve as a proxy for non-pest systems that may also experience rapid environmental change. In order to more accurately understand how adaptation in a population might have occurred, reliable genetic and demographic estimates of population size, mutation rate, and recombination rate are necessary. Therefore, we first estimate these parameters and test for evidence of hard selective sweeps in two rapidly adapting pest populations of the Colorado potato beetle.

Classical expectations for single locus adaptive mutations have been challenged by growing evidence of polygenic resistance traits (Pélissié et al., 2022; Kreiner et al., 2021; Wybouw et al., 2019). However, there are well documented examples of both single locus and polygenic resistance evolution (ffrench-Constant 2013) and the relative importance of these selective regimes might relate to fundamental population genetic properties (Barton 2010; Hermisson and Pennings 2005). Here we provide evidence that hard selective sweeps occur in CPB at unique loci attributed to distinct resistance mechanisms and occur independently in geographically isolated populations. Identifying these independently derived hard sweeps contributes to our understanding of CPB adaptation, along with prior studies that suggest soft selective sweeps are drawn from standing variation across the range of CPB (Crossley et al., 2017; Pélissié et al., 2022). Additionally, through an analysis of demographic history and recombination patterns, we provide an improved understanding of pest population history and sources of genomic variation in CPB, which will improve predictive modeling of resistance evolution in agricultural ecosystems.

### Demographic History of Colorado Potato Beetle Pest Populations

In contrast to introduced pests, which experience a genetic bottleneck prior to invasion (North et al., 2021), native pests are thought to retain much of their genetic variation. Unlike other major agricultural pest species, CPB experienced a host expansion onto potato and switched to pest status in part of its endemic range, North America (eastern Nebraska) in 1859 (Walsh 1866). The host expansion of CPB allowed for the retention of high standing genetic diversity as the pest extended its range eastward (Cohen et al. 2020; Pélissié et al., 2022; Izzo et al., 2018). Geographical structure was shown in earlier population genetic studies using microsatellites, AFLPs, and mitochondrial DNA (Grapputo et al., 2005; Izzo et al., 2018). Our models suggest that CPB pest populations in the Midwest and Eastern U.S. diverged from a founder population of approximately ∼6,700 individuals nearly 160 years ago (∼320 generations) and increased in size in Wisconsin (2.2 fold) and New York (5.5-fold). Contemporary work with whole genome resequencing data, albeit at much smaller population sample sizes, suggests large effective population sizes occur throughout the pest range of CPB (Pélissié et al., 2022).

Our conservative thresholds for SNP ascertainment and our sample sizes are reasonable to reliably determine effective size per population, yet given the low sequencing depth per individual, we might expect possible heterozygous sites to be mistaken as homozygous (Fumagalli 2013). This can influence the SFS by reducing singletons and causing a bias that results in underestimating demography. As the SFS is integral to estimates of demography and recombination (less so in RAiSD, given this metric relies on linkage disequilibrium and variance between swept regions, in addition to the SFS), our results might lead to an underestimate in effective population size (Crawford et al., 2012). We based our analysis of recombination on the dadi estimates, which were ∼4-fold larger than estimates from the Stairway plot analysis. The dadi estimate was obtained on the full joint-SFS after SNP filtering for a global minor allele frequency of 0.05 and under assumptions of a constant size demographic model. In our Stairway plot analysis, we implemented model settings to ignore singletons and recovered a declining population. While there is uncertainty about the true effective size, we note that assuming slightly larger effective size should affect population-based recombination rate (ρ) and not the genome-wide pattern of recombination.

The migration history, contemporary land usage and pest management practices differ significantly between Wisconsin and New York. Historically, potato acreage in Wisconsin was rather dispersed, but covered approximately 3 million acres (∼1200km2) in the early 20th century (Crossley et al., 2019). It subsequently became concentrated in several growing regions comprising ∼75,000 acres (300km2) today. This contrasts with the potato growing history of Long Island, which was very concentrated and encompassed 70,000 acres or ∼300 km2. Present-day estimates suggest potato farming comprises only ∼300 acres (∼1.25km2; Alyokhin et al., 2015). While this dramatic reduction in potato growing acreage (habitat) might account for the steep decline in population size that was estimated to occur 50-100 years ago (**Figure 1**), as well as the slight reduction in the number of SNPs identified within swept regions, this timing is also coincident with the introduction of modern synthetic insecticides. The variability of the lower confidence interval for Long Island might be due to a lack of coalescent events that provide an estimate for effective population size. Resistance history to insecticides varies between Hancock and Long Island, as the Long Island populations developed resistance to every major class of insecticides and have the highest tolerance to novel chemicals of any CPB population (Olson et al., 2000).

There has been some debate about the dynamics of this species’ range expansion, particularly whether it involved admixture among multiple non-pest source populations (Izzo et al., 2018), or a much earlier colonization of the Midwestern U.S. than recorded in the literature (Pélissié et al., 2022). Our estimate of ∼160 years of divergence between Wisconsin and New York populations approximates the historically observed expansion of CPB onto cultivated potato in the 1850s and within a small margin of error for the known CPB occurrence in both states (Tower 1906; Izzo et al., 2018). During the mid-to late-19th century, there was a large continuous band of cultivated potato extending from Nebraska to the Midwest and Eastern U.S. (Crossley et al., 2020), which provided a dispersal corridor for the pest lineage to quickly spread to the Atlantic seaboard by 1874 (Casagrande 1987; Riley 1877). While the effective population size trajectory through time differs between Long Island and Wisconsin, with Long Island having a larger effective population size (supported by both dadi and stairway plot results), there is broad overlap in confidence intervals and a convergence in estimates around the ancestral population size.

In our dadi analysis, a variable population effective size was not favored over a constant population size demographic model, although extant population size was estimated to be several fold larger in both regions. The stairway plot suggested a decline in effective population size for both extant pest populations. The discrepancy between the two results is most likely explained by our methodological choices, as we ignored singletons in the stairway plot. In contrast, we estimated a complete 2D allele frequency spectrum in dadi after filtering for minor allele frequency, but this analysis retained singleton SNPs. While the estimates for demographic history vary, the stairway plot estimates result in large confidence intervals around the effective size mean and this encompasses our estimates from dadi (Mazet et al., 2016). Other factors, such as gene flow from a structured population, could generate a spurious signature of a recent bottleneck. In a previous study, Crossley et al. (2017) showed that Wisconsin populations were weakly structured. However, it is not clear that this accounts for the declining population size trajectory of CPB samples from Hancock. Perhaps ecological factors, such as a declining extent of Wisconsin potato acreage in the late 19th century (Crossley et al., 2021), may explain reductions in effective size since colonization.

### Regional Differences in Recombination

We found similar levels of the per generation recombination rate (r) in both HAN and LI, although the population-based recombination rate (ρ) is higher in LI due to its larger effective population size estimate. Both populations have regions of high and low recombination, with many more low recombining regions. Low recombining regions may be indicative of positive or purifying selection, and these regions intersect genes associated with different modes of insecticide resistance, including target site insensitivity (glutamate receptors XP_023012213 and XP_023012139, which are targets of the abamectin IRAC MOA Group 6) and detoxification (carboxylesterase XP_023018836 and CYP genes including one CYPb-c1, two CYP6a2, three CYP6a23, and five CYP9e2s. CYPs have been repeatedly associated with metabolic detoxification of xenobiotics (Scott and Wen 2001; ffrench-Constant et al., 2004; Bass and Field 2011), resistance to particular insecticidal modes of action (Daborn et al., 2001; Amichot et al., 2004), and cross resistance to multiple modes of action (Peng et al., 2016; Zhu et al., 2016). Our estimates of population-based recombination rates (ρ = 4Ne r) ranged from 0.163 to 0.316, while rates for the X chromosome (ρ = 2Ner) ranged from 0.086 to 0.195. To our knowledge, these are the first recombination rate estimates based on population genomic data for Coleoptera. The rates are similar to estimates from Anopheles (Nelson et al., 2021), but approximately an order of magnitude lower than Drosophila melanogaster (Chan et al., 2012) and several bee species (Jones et al., 2019). However, it is well known that recombination rates can vary widely among closely related species, populations, and even between the sexes (Smukowski and Noor 2011; Kong et al., 2010; Stapley et al., 2017).

### Hard Selective Sweeps and Models of Insecticide Adaptation

Metabolic detoxification has emerged as the most ubiquitous type of resistance in arthropod pests, which may not be surprising given its diversity of molecular pathways and enzyme families, as well as its evolutionary function in protecting arthropods against plant allelochemicals (Li et al., 2007). Detoxifying enzymes are generally classified into three phases based on their relationship to xenobiotic substrates (Li et al., 2007; Heckel 2012). Phase I: functionalization mediated by cytochrome P450s (CYPs); Phase II: conjugation to hydrophilic substrates mediated by glutathione-S-transferases (GSTs) and esterases; and Phase III: excretion mediated by ATP-binding cassettes (ABC transporters). Target site insensitivity, usually involving a point mutation, is another prevalent mechanism of xenobiotic resistance that has been associated with pyrethroid and DDT resistance in *Musca domestica* (Rinkevich et al., 2012; Rinkevich et al., 2006), *Anopheles gambiae* (Ranson et al., 2000) *Culex pipiens* (Martinez-Torres et al., 1999), *L. decemlineata* (Hawthorne 2001), and tobacco budworm, *Heliothis virescens* (Park and Taylor 1997).

Using parameter estimates to generate neutral data under a population growth model for each population, we ascertained probable false positive rates for the μ statistic in RAiSD at cutoffs of 5%, 0.5%, and 0.05%. A growth model was chosen to be conservative, even though our demographic reconstruction indicated a constant size model. No outliers of selection (large μ values) are identified in the neutral simulations (**Figure 3**). In contrast, the observed data shows a clear peak of large μ values for both LI and HAN. We opted to use a 0.05% threshold on our observed SNP data sets to identify the most strongly supported hard sweeps. Both pest populations showed signatures of strong selective sweeps at genes associated with agricultural adaptation. While the genes involved in the selective sweeps were unique to each population, the genes had similar roles in detoxification, host detection, and defense, based on gene ontology (**Supplemental Tables S3-S4**). Of the nine shared genes, two have an association with insecticide resistance: a carboxylesterase (Phase II; XP_023025254) and an ABC transporter (Phase III; XP_023026978), while the other shared genes are associated with neuronal development and function, GMP synthase and metabolism (**Table 1)**. We acknowledge that given the shared demography of these populations, the signatures of selection at these loci could predate historical insecticide use, so further work is needed to clarify the age of selected alleles.

Genes in these protein families have been linked with pesticide detoxification in CPB and other insect pests (Jao and Casida 1974; Bhatt et al., 2020; Cohen et al., 2020; Crossley et al., 2019; Lü et al., 2015). The Long Island population is one of the most abundant and insecticide resistant CPB populations in the literature (Alyokhin et al. 2015; Dively et al., 2020). Tolerance has been attributed to constitutive overexpression of detoxification genes (CYPs, GSTs, and esterases, among others; Clements et al., 2018). However, one of the unique swept loci in the LI population is a major target site for neonicotinoids, the nicotinic acetylcholinesterase receptor subunit. This is not differentially expressed in resistant samples (Dively et al., 2020), despite this population having much higher tolerance for imidacloprid than WI. A CYP locus (CYP9e2) was also swept in the Long Island population and is among other CYPs in shared low-recombination regions with Hancock, WI. For Hancock, WI, sweeps include a known trans-regulatory element for imidacloprid resistance (cap ‘n’ collar), which induces co-expression of ABC transporters, GSTs, CYPs, and esterases (Gaddelapati et al., 2018). The Hancock population also has evidence of selection for a biologically integral defensive enzyme (salicyl alcohol oxidase, LDEC016943), which catalyzes the formation of a volatile deterrent salicylaldehyde in *Chrysomela tremulae* and *C. populi* larvae (Michalski et al., 2008). Its role and relevance in CPB has not been determined.

Relative to other studies that have identified selective sweeps in insect pests, we identify more hard swept regions, yet a similar number of genes within these swept regions (Weedall et al., 2020; Nam et al., 2020; Calla et al., 2020). Until recently, most research on CPB focused on well-known target genes and metabolic pathways (Alyokhin et al. 2008). As next generation sequencing technology and genomic resources have become available for CPB (Kumar et al., 2014; Schoville et al., 2018; Cohen et al., 2020), there has been a rapid expansion of knowledge into the genetic basis of pesticide resistance using genome scans as an agnostic approach to gene discovery. With respect to prior comparative population genetic studies of CPB, which had presented extensive candidate gene lists (>8,700 genes from a more inclusive demographic sampling; Pélissié et al., 2022), this work suggests a small set of genes (<150 per population) undergo hard selective sweeps. Compared to Pélissié *et al*. (2022), we find that ABC transporters and an esterase, swept in both NY and WI, were also significant in their genome scan results. Additionally, 13 and 17 genes from their PCAdapt analysis are shared in our analysis of Wisconsin and New York samples, respectively (**Supplemental Table S3**). The size of the swept regions around significant SNPs is larger than the average size of all regions around all SNPs.

The distribution and number of SNPs between these populations suggests that they are distinct haplotypes that have been selected independently in each region. Together, these results imply a combined effect of newly determined hard and previously described soft selected sweeps on CPB genes that support agricultural adaptation maintained by large population size and standing genetic diversity.

### Leveraging Population Genomics for Integrated Pest Management

The ability of CPB to rapidly evolve resistance continues to challenge the sustainability of potato production (Alyokhin et al. 2015), which is an economically important vegetable crop with an annual production valued at $3.74 billion (USDA 2017). Due to a general lack of effective natural enemies of CPB, potato production has historically depended upon insecticides to control the beetle (Hare 1990). Without a clear understanding of *how* CPB and other successful pests continue to evolve resistance, resistance will continue to develop in a matter of generations and growers will lack effective strategies to manage CPB over the long-term.

Interestingly, we determine similar selective regimes in divergent populations despite dramatically different management practices and measured insecticide tolerances. Expanding baseline knowledge of the genetic loci and gene networks involved in resistance can help mitigate the spread of resistance into susceptible populations by allowing for biomonitoring. However, a potentially more important outcome of this work is to enable a refinement of IPM strategies for the future use of novel pesticidal chemistries. Improved understanding of population genetic parameters, including their variation among geographical regions, allows for more effective predictive modeling of resistance evolution (Karlsson et al., 2020). Drawing on examples of insecticide, fungicide, and antibiotic resistance, Beckie et al. (2021) have argued that efforts to minimize selection and impede dispersal in taxa that exhibit cross-resistance may need to be tailored to specific pest species for improved IPM outcomes. Due to substantial standing genetic variation and a tendency to locally evolve resistance by both hard and soft sweeps, management of CPB should leverage modeling approaches that examine not only single large effect *de novo* mutations, but also accumulation of polygenic resistance traits from background genetic diversity (Haridas et al., 2018) in discrete regional pest populations.

## Acknowledgements

We thank Sandra Menasha and Michael Crossley for providing the beetle samples used in this study. This work was supported by a USDA NIFA AFRI Exploratory Grant (2015-67030-23495), a USDA-National Potato Council award (58-5090-7-073), and a Hatch Award (WIS02004). In addition, Z. Cohen was supported by the Wisconsin Potato and Vegetable Growers Association in the form of a Wisconsin Distinguished Graduate Fellowship Award (#233JJ17).

## Supplementary Material

**Supplemental File 1**. Contains Supplemental Tables S1-S6, Supplemental Figures S1-S8, and Supplemental Methods (.docx format).

## Data Accessibility Statement

Raw genomic read data for each population have been deposited on NCBI (PRJNA753140).

## References

Akaike, H. (1973). Information theory and an extension of the maximum likelihood principle. In B. N. Petrov & F. Csáki (Eds.), 2nd international symposium on infor-mation theory(pp. 267– 281). Budapest, Hungary: Akadémia Kiadó.

Alachiotis N, Pavlidis P. 2018. RAiSD detects positive selection based on multiple signatures of a selective sweep and SNP vectors. Commun Biol. 1(1). doi:10.1038/s42003-018-0085-8.

Alyokhin A, Baker M, Mota-Sanchez D, Dively G, Grafius E. 2008. Colorado potato beetle resistance to insecticides. Am J Potato Res. 85(6):395–413. doi:10.1007/s12230-008-9052-0.

Alyokhin A, Mota-Sanchez D, Baker M, Snyder WE, Menasha S, Whalon M, Dively G, Moarsi WF. 2015. The Red Queen in a potato field: Integrated pest management versus chemical dependency in Colorado potato beetle control. Pest Manag Sci. 71(3):343–356. doi:10.1002/ps.3826.

Amichot M, Tarés S, Brun-Barale A, Arthaud L, Bride JM, Bergé JB. 2004. Point mutations associated with insecticide resistance in the Drosophila cytochrome P450 Cyp6a2 enable DDT metabolism. Eur J Biochem. 271(7):1250–1257. doi:10.1111/j.1432-1033.2004.04025.x.

Barton N. 2010. Understanding adaptation in large populations. PLoS Genet. 6(6):1–3. doi:10.1371/journal.pgen.1000987.

Bass C, Field LM. 2011. Gene amplification and insecticide resistance. Pest Manag Sci. 67(8):886–890. doi:10.1002/ps.2189.

Bass, C., Denholm, I., Williamson, M.S., Nauen, R., 2015. The global status of insect resistance to neonicotinoid insecticides. Pestic. Biochem. Physiol. 121, 78–87.

Beckie HJ, Busi R, Lopez-Ruiz FJ, Umina PA. 2021. Herbicide resistance management strategies: how do they compare with those for insecticides, fungicides and antibiotics? Pest Manag Sci. 77(7):3049–3056. doi:10.1002/ps.6395.

Bhaskara S, Dean ED, Lam V, Ganguly R. 2006. Induction of two cytochrome P450 genes, Cyp6a2 and Cyp6a8, of Drosophila melanogaster by caffeine in adult flies and in cell culture. Gene. 377(1–2):56–64. doi:10.1016/j.gene.2006.02.032.

Bhatt P, Bhatt K, Huang Y, Lin Z, Chen S. 2020. Esterase is a powerful tool for the biodegradation of pyrethroid insecticides. Chemosphere. 244:125507. doi:10.1016/j.chemosphere.2019.125507.

Boiteau G, Alyokhin A, Ferro D. 2003. The Colorado potato beetle in movement. The Canadian Entomologist. 135(1)1–22.

Brévault, T., Heuberger, S., Zhang, M., Ellers-Kirk, C., Ni, X., Masson, L., Li, X., Tabashnik, B.E., Carrière, Y., 2013. Potential shortfall of pyramided transgenic cotton for insect resistance management, 110, pp. 5806–5811.

Brevik K, Schoville SD, Mota-Sanchez D, Chen YH. 2018. Pesticide durability and the evolution of resistance: A novel application of survival analysis. Pest Manag Sci.: 1–27. doi:10.1002/ps.4899.

Calla B, Demkovich M, Siegel JP, Viana JPG, Walden KKO, Robertson HM, Berenbaum MR. 2021. Selective Sweeps in a Nutshell: The Genomic Footprint of Rapid Insecticide Resistance Evolution in the Almond Agroecosystem. Genome Biol Evol. 13(1):1–14. doi:10.1093/gbe/evaa234.

Chan AH, Jenkins PA, Song YS. 2012. Genome-wide fine-scale recombination rate variation in Drosophila melanogaster. PLoS Genet. 8(12). doi:10.1371/journal.pgen.1003090.

Chen H, Patterson N, Reich D. 2010. Population differentiation as a test for selective sweeps. Genome Res. 20(3):393–402. doi:10.1101/gr.100545.109.

Clements J, Sanchez-Sedillo B, Bradfield CA, Groves RL. 2018. Transcriptomic analysis reveals similarities in genetic activation of detoxification mechanisms resulting from imidacloprid and chlorothalonil exposure. PLoS One. 13(10):1–14. doi:10.1371/journal.pone.0205881.

Clements J, Schoville S, Peterson N, Lan Q, Groves RL. 2016. Characterizing Molecular Mechanisms of Imidacloprid Resistance in Select Populations of Leptinotarsa decemlineata in the Central Sands Region of Wisconsin. PLoS One. 11(1):e0147844. doi:10.1371/journal.pone.0147844.

Cohen ZP, Brevik K, Chen YH, Hawthorne DJ, Weibel BD, Schoville SD. 2020 Oct 23. Elevated rates of positive selection drive the evolution of pestiferousness in the Colorado potato beetle (Leptinotarsa decemlineata, Say). Mol Ecol. doi:10.1111/mec.15703.

Cohen ZP, Hawthorne DJ, Schoville SD. 2021. The role of structural variants in pest adaptation and genome evolution of the Colorado potato beetle, Leptinotarsa decemlineata (Say). Authorea. doi: 10.22541/au.163109527.73747035/v1

Crawford JE, Lazzaro BP. 2012. Assessing the accuracy and power of population genetic inference from low-pass next-generation sequencing data. Front Genet. 3(APR):1–13. doi:10.3389/fgene.2012.00066.

Crossley MS, Chen YH, Groves RL, Schoville SD. 2017. Landscape genomics of Colorado potato beetle provides evidence of polygenic adaptation to insecticides. Molecular

Crossley MS, Rondon SI, Schoville SD. 2019. Effects of contemporary agricultural land cover on Colorado potato beetle genetic differentiation in the Columbia Basin and Central Sands. (April):1–10. doi:10.1002/ece3.5489.

Daborn P, Boundy S, Yen J, Pittendrigh B, Ffrench-Constant R. 2001. DDT resistance in Drosophila correlates with Cyp6g1 over-expression and confers cross-resistance to the neonicotinoid imidacloprid. Mol Genet Genomics. 266(4):556–563. doi:10.1007/s004380100531.

De-Kayne R, Frei D, Greenway R, Mendes SL, Retel C, Feulner PGD. 2021. Sequencing platform shifts provide opportunities but pose challenges for combining genomic data sets. Mol Ecol Resour. 21(3):653–660. doi:10.1111/1755-0998.13309.

Dively, G. P., Crossley, M. S., Schoville, S. D., Steinhauer, N., & Hawthorne, D. J. (2020). Regional differences in gene regulation may underlie patterns of sensitivity to novel Insecticides in Leptinotarsa decemlineata. Pest Management Science, 12, 4278–4285 Ecology. 26(22):6284–6300

Ferro, D. N., J. A. Logan, R. H. Voss, and J. S. Elkinton. “Colorado potato beetle (Coleoptera: Chrysomelidae) temperature-dependent growth and feeding rates.” Environmental entomology 14, no. 3 (1985): 343-348.

ffrench-Constant RH, Daborn PJ, Le Goff G. 2004. The genetics and genomics of insecticide resistance. Trends Genet. 20(3):163–170. doi:10.1016/j.tig.2004.01.003.

ffrench-Constant RH. 2013. The molecular genetics of insecticide resistance. Genetics. 194(4):807–815. doi:10.1534/genetics.112.141895.

Forgash J. 1985. Insecticide resistance in the Colorado potato beetle. In: Proceedings of the Symposium on the Colorado Potato Beetle, XVIIth International Congress of Entomology. p. 33–52.

Fumagalli M. 2013. Assessing the effect of sequencing depth and sample size in population genetics inferences. PLoS One. 8(11):e79667. doi:10.1371/journal.pone.0079667.

Gaddelapati SC, Kalsi M, Roy A, Palli SR. 2018. Cap “n” collar C regulates genes responsible for imidacloprid resistance in the Colorado potato beetle, Leptinotarsa decemlineata. Insect Biochem Mol Biol. 99:54–62. doi:10.1016/J.IBMB.2018.05.006.

Georghiou GP, Taylor CE (1977) Genetic and biological influences in the evolution of insecticide resistance. Journal of Economic Entomology 70: 319–323. https://doi.org/10.1093/jee/70.3.319

Gould, F., Brown, Z. S., & Kuzma, J. (2018). Wicked evolution: Can we address the sociobiological dilemma of pesticide resistance? Science, 360(6390), 728–732. https://doi.org/10.1126/scien ce.aar3780

Grapputo A, Boman S, Lindström L, Lyytinen A, Mappes J. 2005. The voyage of an invasive species across continents: Genetic diversity of North American and European Colorado potato beetle populations. Mol Ecol. 14(14):4207–4219. doi:10.1111/j.1365-294X.2005.02740.x.

Groeters, F. R., and B. E. Tabashnik, 2000 Roles of selection intensity, major genes, and minor genes in evolution of insecticide resistance. J. Econ. Entomol. 93: 1580–1587.

Gutenkunst RN, Hernandez RD, Williamson SH, Bustamante CD. 2009. Inferring the joint demographic history of multiple populations from multidimensional SNP frequency data. PLoS Genet. 5(10). doi:10.1371/journal.pgen.1000695.

Haridas C V., Tenhumberg B. 2018. Modeling effects of ecological factors on evolution of polygenic pesticide resistance. J Theor Biol. 456:224–232. doi:10.1016/j.jtbi.2018.07.034.

Hartl, Daniel L., and Andrew G. Clark. Principles of population genetics. Vol. 116. Sunderland, MA: Sinauer associates, 1997.

Hawkins NJ, Bass C, Dixon A, Neve P. 2019. The evolutionary origins of pesticide resistance. Biol Rev. 94(1):135–155. doi:10.1111/brv.12440.

Hawthorne DJ. 2001. AFLP-based genetic linkage map of the Colorado potato beetle Leptinotarsa decemlineata: sex chromosomes and a pyrethroid-resistance candidate gene. Genetics. 158(2):695–700.

Hawthorne DJ. 2003. Quantitative trait locus mapping of pyrethroid resistance in Colorado potato beetle, Leptinotarsa decemlineata (Say) (Coleoptera: Chrysomelidae). J Econ Entomol. 96(4):1021–1030.

Heckel, D. G., Gahan, L. J., Baxter, S. W., Zhao, J. Z., Shelton, A. M., Gould, F., & Tabashnik, B. E. (2007). The diversity of Bt resistance genes in species of Lepidoptera. Journal of Invertebrate Pathology, 95(3), 192–197. https://doi.org/10.1016/j.jip.2007.03.008

Hermisson J, Pennings PS. 2005. Soft sweeps: Molecular population genetics of adaptation from standing genetic variation. Genetics. 169(4):2335–2352. doi:10.1534/genetics.104.036947.

Hoy MA. 1998. Myths, models and mitigation of resistance to pesticides. Philos Trans R Soc London B Biol Sci. 353:1787–95.

Hudson RR. 2002. Generating samples under a Wright-Fisher neutral model. Bioinformatics 18: 337–338. doi:10.1093/bioinformatics/18.2.337.

Huseth AS, Groves RL, Chapman SA, Alyokhin A, Kuhar TP, Macrae I V, Szendrei Z, Nault BA. 2014. Managing Colorado Potato beetle insecticide resistance: new tools and strategies for the next decade of pest control in potato. J Integrated Pest Management. 5(4):1–8. doi:10.1603/IPM14009.

Ioannidis PM, Grafius E, Whalon ME. 1991. Patterns of insecticide resistance to azinphosmethyl, carbofuran, and permethrin in the Colorado potato beetle (Coleoptera: Chrysomelidae). J Econ Entomol. 84:1417–1423.

IRAC. 2016. Insecticide resistance action committee database. http://www.irac-online.org/.

Izzo VM, Chen YH, Schoville SD, Wang C, Hawthorne DJ. 2018. Origin of Pest Lineages of the Colorado Potato Beetle (Coleoptera: Chrysomelidae). J Econ Entomol. 1859:1–11. doi:10.1093/jee/tox367. http://academic.oup.com/jee/advance-article/doi/10.1093/jee/tox367/4818462.

Jacques RL. 1988. The potato beetles: the genus Leptinotarsa in North America (Coleoptera, Chrysomelidae). Leiden; New York: E.J. Brill.

Jao LT, Casida JE. 1974. Insect Pyrthroid-Hydrolyzing Esterases. Pestic Biochem Physiol. 4:465–472.

Jolivet P, Hsiao TH, Petitpierre E. 1988. Biology of Chrysomelidae. http://link.springer.com/10.1007/978-94-009-3105-3.

Jones JC, Wallberg A, Christmas MJ, Kapheim KM, Webster MT, Singh N. 2019. Extreme Differences in Recombination Rate between the Genomes of a Solitary and a Social Bee. Mol Biol Evol. 36(10):2277–2291. doi:10.1093/molbev/msz130.

Karlsson Green K, Stenberg JA, Lankinen Å. 2020. Making sense of Integrated Pest Management (IPM) in the light of evolution. Evol Appl. 13(8):1791–1805. doi:10.1111/eva.13067.

Kong A, Thorleifsson G, Gudbjartsson DF, Masson G, Sigurdsson A, Jonasdottir Aslaug, Walters GB, Jonasdottir Adalbjorg, Gylfason A, Kristinsson KT, et al. 2010. Fine-scale recombination rate differences between sexes, populations and individuals. Nature. 467(7319):1099–1103. doi:10.1038/nature09525.

Korneliussen TS, Albrechtsen A, Nielsen R. 2014. ANGSD: Analysis of Next Generation Sequencing Data. BMC Bioinformatics. 15:356. doi: 10.1186/s12859-014-0356-4.

Kreiner JM, Tranel PJ, Weigel D, Stinchcombe JR, Wright SI. 2021. The genetic architecture and population genomic signatures of glyphosate resistance in Amaranthus tuberculatus. Mol Ecol. 30(21): 1–17. doi:10.1111/mec.15920.

Kumar A, Congiu L, Lindström L, Piiroinen S, Vidotto M, Grapputo A. 2014. Sequencing, de Novo assembly and annotation of the colorado potato beetle, Leptinotarsa decemlineata, transcriptome. PLoS One. 9(1). doi:10.1371/journal.pone.0086012.

Kuznetsova I, Lugmayr A, Siira SJ, Rackham O, Filipovska A. 2019. CirGO: An alternative circular way of visualizing gene ontology terms. BMC Bioinformatics. 20(1):1–7. doi:10.1186/s12859-019-2671-2.

Li H, Handsaker B, Wysoker A, Fennell T, Ruan J, Homer N, Marth G, Abecasis G, Durbin R, others. 2009. The sequence alignment/map format and SAMtools. Bioinformatics. 25(16): 2078–2079.

Li X, Schuler MA, Berenbaum MR. 2007. Molecular Mechanisms of Metabolic Resistance to Synthetic and Natural Xenobiotics. Annu Rev Entomol. 52(1):231–253. doi:10.1146/annurev.ento.51.110104.151104.

Li, H., 2013 Aligning sequence reads, clone sequences and assembly contigs with BWA-MEM. arXiv: 1303.3997.

Lin G, Chai J, Yuan S, Mai C, Cai L, Murphy RW, Zhou W, Luo J. 2016. Vennpainter: A tool for the comparison and identification of candidate genes based on Venn diagrams. PLoS One. 11(4): 5–8. doi:10.1371/journal.pone.0154315.

Liu X, Fu YX. 2020. Stairway Plot 2: demographic history inference with folded SNP frequency spectra. Genome Biol. 21(1):1–9. doi:10.1186/s13059-020-02196-9.

Lohmueller, Kirk E., Jeremiah D. Degenhardt, and Alon Keinan. Sex-averaged recombination and mutation rates on the X chromosome: a comment on Labuda et al. Amer J Human Gen. 86(6): 978

Lü FG, Fu KY, Li Q, Guo WC, Ahmat T, Li GQ. 2015. Identification of carboxylesterase genes and their expression profiles in the Colorado potato beetle Leptinotarsa decemlineata treated with fipronil and cyhalothrin. Pestic Biochem Physiol. 122:86–95. doi:10.1016/j.pestbp.2014.12.015.

Mamidala P, Jones SC, Mittapalli O. 2011. Metabolic Resistance in Bed Bugs. Insects. 2(1): 36–48. doi:10.3390/insects2010036

Martinez-Torres D, Chevillon C, Brun-Barale A, Berge JB, Pasteur N, Pauron D. 1999. Voltage-dependent Na+ channels in pyrethroid-resistant Culex pipiens L mosquitoes. Pestic Sci. 55(10):1012–1020. doi:10.1002/(SICI)1096-9063(199910)55:10<1012::AID-PS39>3.0.CO;2-5.

Mazet O, Rodríguez W, Grusea S, Boitard S, Chikhi L. 2016. On the importance of being structured: Instantaneous coalescence rates and human evolution-lessons for ancestral population size inference? Heredity 116(4):362–371. doi:10.1038/hdy.2015.104.

McDougall P. 2011. AgriService Report. Phillips McDougall, Midlothian.

Michalski C, Mohagheghi H, Nimtz M, Pasteels J, Ober D. 2008. Salicyl alcohol oxidase of the chemical defense secretion of two chrysomelid leaf beetles: Molecular and functional characterization of two new members of the glucose-methanol-choline oxidoreductase gene family. J Biol Chem. 283(28):19219–19228. doi:10.1074/jbc.M802236200.

Nam K, Nhim S, Robin S, Bretaudeau A, Nègre N, d’Alençon E. 2020. Positive selection alone is sufficient for whole genome differentiation at the early stage of speciation process in the fall armyworm. BMC Evol Biol. 20(1):1–16. doi:10.1186/s12862-020-01715-3.

Nelson JT, Cornejo OE, Consortium A. 2021 Jan 1. Evolutionary implications of recombination differences across diverging populations of Anopheles

North HL, McGaughran A, Jiggins CD. 2021. Insights into invasive species from whole-genome resequencing. Mol Ecol.(April):1–20. doi:10.1111/mec.15999.

O’Reilly PF, Birney E, Balding DJ. 2008. Confounding between recombination and selection, and the Ped/Pop method for detecting selection. Genome Res. 18(8):1304–1313. doi:10.1101/gr.067181.107.

Olson ER, Dively GP, Nelson JO. 2000. Baseline Susceptibility to Imidacloprid and Cross Resistance Patterns in Colorado Potato Beetle (Coleoptera: Chrysomelidae) Populations. J Econ Entomol. 93(2):447–458. doi:10.1603/0022-0493-93.2.447.

Oppold A-M, Pfenninger M: Direct estimation of the spontaneous mutation rate by short-term mutation accumulation lines in Chironomus riparius. Evolution Letters 2017, 1–2:86–92

Park Y, Taylor MFJ. 1997. A novel mutation L1029H in sodium channel gene hscp associated with pyrethroid resistance for Heliothis virescens (lepidoptera: noctuidae). Insect Biochem Mol Biol. 27(1):9–13. doi:10.1016/S0965-1748(96)00077-X.

Pélissié B, Chen YH, Cohen ZP, Crossley MS, Hawthorne DJ, Izzo V, Schoville SD. 2022. Genome resequencing reveals rapid, repeated evolution in the Colorado potato beetle. Mol Biol Evol 39(2): msac016.

Pélissié B, Crossley MS, Cohen ZP, Schoville SD. 2018. Rapid evolution in insect pests: the importance of space and time in population genomics studies. Curr Opin Insect Sci. 26. doi:10.1016/j.cois.2017.12.008.

Pelletier Y, Horgan FG, Pompon J. 2011. Potato resistance to insects. Am J Plant Sci Biotechnol. 5:37–52.

Peng T, Pan Y, Yang C, Gao X, Xi J, Wu Y, Huang X, Zhu E, Xin X, Zhan C, et al. 2016. Over-expression of CYP6A2 is associated with spirotetramat resistance and cross-resistance in the resistant strain of Aphis gossypii Glover. Pestic Biochem Physiol. 126:64–69. doi:10.1016/j.pestbp.2015.07.008.

Portik DM, Leaché AD, Rivera D, Barej MF, Burger M, Hirschfeld M, Rödel MO, Blackburn DC, Fujita MK. 2017. Evaluating mechanisms of diversification in a Guineo-Congolian tropical forest frog using demographic model selection. Mol Ecol. 26(19):5245–5263. doi:10.1111/mec.14266.

Quinlan AR, Hall IM. 2010. BEDTools: A flexible suite of utilities for comparing genomic features. Bioinformatics. 26(6):841–842. doi:10.1093/bioinformatics/btq033.

R Core Team (2021). R: A language and environment for statistical computing. R Foundation for Statistical Computing, Vienna, Austria. URL https://www.R-project.org/.

Ranson H, Jensen B, Vulule JM, Wang X, Hemingway J, Collins FH. 2000. Identification of a point mutation in the voltage-gated sodium channel gene of Kenyan Anopheles gambiae associated with resistance to DDT and pyrethroids. Insect Mol Biol. 9(5):491–497. doi:10.1046/j.1365-2583.2000.00209.x.

Riley, C. V. 1877. The Colorado potato beetle: with suggestions for its repression and methods of destruction. Routledge, London

Rinkevich F.D., Zhang L, Hamm RL, Brady S.G., Lazzaro B.P., Scott J.G., Frequencies of the pyrethroid resistance alleles of Vssc1 and CYP6D1 in house flies from the eastern United States, Insect Mol. Biol. 15 (2006) 157–167.

Rinkevich FD, Su C, Lazo TA, Hawthorne DJ, Tingey WM, Naimov S, Scott JG. 2012. Multiple evolutionary origins of knockdown resistance (kdr) in pyrethroid-resistant Colorado potato beetle, Leptinotarsa decemlineata. Pestic Biochem Physiol. 104(3):192–200.

Roush, R. T. 1989. Designing resistance management programs: How can you choose? Pestic. Sci. 26: 423–441

Schoville SD, Chen YH, Andersson MN, Benoit JB, Bowsher JH, Brevik K, Cappelle K, Chen MM, Anna K, Childers C, et al. 2017. A model species for agricultural pest genomics : the genome of the Colorado potato beetle, Leptinotarsa decemlineata (Coleoptera : Chrysomelidae)

Scott JG, Wen Z. 2001. Cytochromes P450 of insects: The tip of the iceberg. Pest Manag Sci. 57(10):958–967. doi:10.1002/ps.354.

Scott, J. G., Liu, N., & Wen, Z. (1998). Insect cytochromes P450: Diversity, insecticide resistance and tolerance to plant toxins. Comparative Biochemistry and Physiology Part C, 121, 147–155.

Sharma S., Kooner R., Arora R. (2017) Insect Pests and Crop Losses. In: Arora R., Sandhu S. (eds) Breeding Insect Resistant Crops for Sustainable Agriculture. Springer, Singapore. https://doi.org/10.1007/978-981-10-6056-4_2

Smukowski CS, Noor MAF. 2011. Recombination rate variation in closely related species. Heredity (Edinb). 107(6):496–508. doi:10.1038/hdy.2011.44.

Spence JP, Song YS. 2019. Inference and analysis of population-specific fine-scale recombination maps across 26 diverse human populations. Sci Adv. 5(October). doi:10.1101/532168.

Stapley J, Feulner PGD, Johnston SE, Santure AW, Smadja CM. 2017. Variation in recombination frequency and distribution across eukaryotes: Patterns and processes. Philos Trans R Soc B Biol Sci. 372(1736). doi:10.1098/rstb.2016.0455.

Supek F, BoŠnjak M, Škunca N, Šmuc T. 2011. Revigo summarizes and visualizes long lists of gene ontology terms. PLoS One. 6(7). doi:10.1371/journal.pone.0021800.

Supek F, Bošnjak M, Škunca N, Šmuc T. 2011. Revigo summarizes and visualizes long lists of gene ontology terms. PloS One. 6(7). doi:10.1371/journal.pone.0021800.

Thomas GWC, Dohmen E, Hughes DST, Murali SC, Poelchau M, Glastad K, Anstead CA, Ayoub NA, Batterham P, Bellair M, et al. 2020. Gene content evolution in the arthropods. Genome Biol. 21(1):1–14. doi:10.1186/s13059-019-1925-7.

Tower WL. 1906. An Investigation of evolution in chrysomelid beetles of the genus Leptinotarsa.

Walsh BD. 1866. The new potato bug. Pract Entomol. 2:13–16.

Weedall GD, Riveron JM, Hearn J, Irving H, Kamdem C, Fouet C, White BJ, Wondji CS. 2020. An Africa-wide genomic evolution of insecticide resistance in the malaria vector Anopheles funestus involves selective sweeps, copy number variations, gene conversion and transposons. PLoS Genet. 16(6):1–29. doi:10.1371/journal.pgen.1008822.

Whalon, M. E., & Mota-Sanchez, D. (2016). Arthropod pesticide resistance database. East Lansing, MI: Michigan State University. http://www.pesticideresistance.org/

Wright, S. 1984. Evolution and the Genetics of Populations, Volume 2: Theory of gene frequencies. University of Chicago press, Chicago.

Wybouw N, Kosterlitz O, Kurlovs AH, Bajda S, Greenhalgh R, Snoeck S, Bui H, Bryon A, Dermauw W, Van Leeuwen T, et al. 2019. Long-term population studies uncover the genome structure and genetic basis of xenobiotic and host plant adaptation in the herbivore tetranychus urticae. Genetics. 211(4):1409–1427. doi:10.1534/genetics.118.301803.

Yeaman S. 2022. Evolution of polygenic traits under global vs local adaptation. Genetics. 220(1). doi:10.1093/genetics/iyab134.

Zhu F, Moural TW, Nelson DR, Palli SR. 2016. A specialist herbivore pest adaptation to xenobiotics through up-regulation of multiple Cytochrome P450s. Sci Rep. 6(August 2015):20421. doi:10.1038/srep20421. http://www.nature.com/articles/srep20421.

