## Supplementary material for "Evidence of hard-selective sweeps suggest independent adaptation to insecticides in Colorado potato beetle (Coleoptera: *Chrysomelidae*) populations": Supplemental_sweep_biorxiv.pdf

**Supplemental File 1**

**Supplementary Methods.** Commands used to conduct genomic data analyses.

#SNP generation for RAiSD and Pyrho:
#Wisconsin:

```
$./angsd/angsd -nThreads 28 -nQueueSize 50 -dobcf 1 -doMaf 1 -dopost 1 -dosaf 1 -gl 2 --ignore-RG 0 -dogeno 1 -anc F_Kansas_60.fasta -doGlf 2 -doMajorMinor 1 -doCounts 1 -remove_bads 1 -minMapQ 30 -minQ 20 -minMaf 0.05 -setMinDepthInd 3 -setMinDepth 84 -setMaxDepth 280 -skipTriallelic 1 -SNP_pval 1e-6 -b WI_bams_update.list -out WI_28_64_snps
```

#New York:

```
$./angsd/angsd -nThreads 28 -nQueueSize 50 -dobcf 1 -doMaf 1 -dopost 1 -dosaf 1 -gl 2 --ignore-RG 0 -dogeno 1 -anc F_Kansas_60.fasta -doGlf 2 -doMajorMinor 1 -doCounts 1 -remove_bads 1 -minMapQ 30 -minQ 20 -minMaf 0.05 -setMinDepthInd 3 -setMinDepth 75 -setMaxDepth 250 -skipTriallelic 1 -SNP_pval 1e-6 -b LI_bam.list -out LI_25_64_snps
```

#stairway plot folded SFS:
### process repeated

```
$./angsd/angsd -b LI_bam.list -gl 2 -anc F_Kansas_60.fasta -ref F_Kansas_60.fasta -rf rf_agenic_regions_FKS.list -dosaf 1 -doCounts 1 -setMinDepthInd 5 -setMinDepth 125 -setMaxDepth 2500 -baq 1 -C 50 -minMapQ 30 -minQ 20 -P 20 -out LI_25_69 -fold 1
```

### dadi joint 2D-SFS:

```
$./angsd/angsd -nThreads 28 -nQueueSize 50 -dobcf 1 -doMaf 1 -dopost 1 -dosaf 1 -gl 2 --ignore-RG 0 -dogeno 1 -anc F_Kansas_60.fasta -doGlf 2 -doMajorMinor 1 -doCounts 1 -remove_bads 1 -minMapQ 30 -minQ 20 -minMaf 0.05 -setMinDepthInd 3 -setMinDepth 84 -setMaxDepth 2800 -skipTriallelic 1 -SNP_pval 1e-6 -b WI_LI_update_bams.list -out WI_LI_611_snps
```

### Allele frequency sampling between reads from different sequencing platforms:
### process repeated

```
$for j in {2..4}; do vcftools --keep shuf"$j".wi.list --vcf nofield_WI_28_64_snps.vcf --freq --out shuf"$j"_WI; done

$for j in `ls new*WI*af`; do awk '{sum += $1} END {print sum/NR}' "$j"; done
```

**Evidence of hard-selective sweeps suggest independent adaptation to insecticides in Colorado potato beetle (Coleoptera: *Chrysomelidae*) populations**

#MS neutral simulation:
### Long Island, NY

$ms 50 100 -t 16.80 -F 5-r 22000.00 50000.00 -eN 0.002 0.1675

#Hancock, WI

$ms 50 100 -t 6.30 -F 5 -r 5700.00 50000.00 -eN 0.0054 0.447

**Evidence of hard-selective sweeps suggest independent adaptation to insecticides in Colorado potato beetle (Coleoptera: *Chrysomelidae*) populations**

#RAiSD neutral thresholds from ms output:
5% ~ LI: 1.68e-10; WI: 1.68e-10
0.5% ~ LI: 2.01e -10; WI: 2.03e-10
0.05% ~ LI: 2.5e-10; WI: 2.4e-10

**Supplementary Table S1.** Samples examined in this study. Genome-wide coverage for museum samples is calculated on each bam file (Pélissié et al., 2022).

| SampleID | SiteType (ref) | Genome-wide Coverage | Year | Latitude | Longitude |
|---|---|---|---|---|---|
| CPB2015-378C | NY | 3.67 | 2015 | 40.905657 | -72.752664 |
| CPB2015-378D | NY | 3.59 | 2015 | 40.905657 | -72.752664 |
| CPB2015-378E | NY | 3.86 | 2015 | 40.905657 | -72.752664 |
| CPB2015-378F | NY | 4.24 | 2015 | 40.905657 | -72.752664 |
| CPB2015-406C | NY | 3.62 | 2015 | 40.905657 | -72.752664 |
| CPB2015-416A | NY | 3.88 | 2015 | 40.905657 | -72.752664 |
| CPB2015-416B | NY | 4.26 | 2015 | 40.905657 | -72.752664 |
| CPB2015-416C | NY | 3.99 | 2015 | 40.905657 | -72.752664 |
| CPB2015-416D | NY | 4.12 | 2015 | 40.905657 | -72.752664 |
| CPB2015-416E | NY | 4.38 | 2015 | 40.905657 | -72.752664 |
| CPB2015-416F | NY | 3.45 | 2015 | 40.905657 | -72.752664 |
| CPB2015-419D | NY | 3.96 | 2015 | 40.905657 | -72.752664 |
| CPB2015-419E | NY | 4.26 | 2015 | 40.905657 | -72.752664 |
| CPB2015-419F | NY | 4.45 | 2015 | 40.905657 | -72.752664 |
| CPB2015-419G | NY | 4.36 | 2015 | 40.905657 | -72.752664 |
| CPB2015-419H | NY | 3.85 | 2015 | 40.905657 | -72.752664 |
| CPB2015-419I | NY | 3.76 | 2015 | 40.905657 | -72.752664 |
| CPB2015-419J | NY | 3.73 | 2015 | 40.905657 | -72.752664 |
| CPB2015-490A | NY | 2.21 | 2015 | 40.905657 | -72.752664 |
| CPB2015-490B | NY | 1.45 | 2015 | 40.905657 | -72.752664 |
| CPBWGS_84 | NY (2022) | 5.03 | 2015 | 40.905657 | -72.752664 |
| CPBWGS_85 | NY (2022) | 4.61 | 2015 | 40.905657 | -72.752664 |
| CPBWGS_86 | NY (2022) | 4.34 | 2015 | 40.905657 | -72.752664 |
| CPBWGS_87 | NY (2022) | 4.95 | 2015 | 40.905657 | -72.752664 |
| CPBWGS_88 | NY (2022) | 3.92 | 2015 | 40.905657 | -72.752664 |
| CPB2015-442A | WI | 4.48 | 2015 | 44.119753 | -89.535683 |
| CPB2015-442B | WI | 4.11 | 2015 | 44.119753 | -89.535683 |
| CPB2015-442C | WI | 5.03 | 2015 | 44.119753 | -89.535683 |
| CPB2015-442D | WI | 4.92 | 2015 | 44.119753 | -89.535683 |
| CPB2015-442E | WI | 4.58 | 2015 | 44.119753 | -89.535683 |
| CPB2015-461 | WI | 3.81 | 2015 | 44.119753 | -89.535683 |
| CPB2015-209A | WI | 4.90 | 2015 | 44.119753 | -89.535683 |
| CPB2015-209B | WI | 4.13 | 2015 | 44.119753 | -89.535683 |
| CPB2015-241A | WI | 4.73 | 2015 | 44.119753 | -89.535683 |
| CPB2015-241B | WI | 4.01 | 2015 | 44.119753 | -89.535683 |
| CPB2015-241C | WI | 4.04 | 2015 | 44.119753 | -89.535683 |
| CPB2015-348A | WI | 4.51 | 2015 | 44.119753 | -89.535683 |
| CPB2015-348B | WI | 4.01 | 2015 | 44.119753 | -89.535683 |

| | | | | | |
|---|---|---|---|---|---|
| CPB2015-348E | WI | 4.62 | 2015 | 44.119753 | -89.535683 |
| CPB2015-499A | WI | 4.01 | 2015 | 44.119753 | -89.535683 |
| CPB2015-499B | WI | 4.14 | 2015 | 44.119753 | -89.535683 |
| CPB2015-348C | WI | 3.74 | 2015 | 44.119753 | -89.535683 |
| CPB2015-348D | WI | 3.78 | 2015 | 44.119753 | -89.535683 |
| CPB2015-348F | WI | 3.75 | 2015 | 44.119753 | -89.535683 |
| CPB2015-417 | WI | 0.07 | 2015 | 44.119753 | -89.535683 |
| CPB2015-504A | WI | 0.004 | 2015 | 44.119753 | -89.535683 |
| CPB2015-504B | WI | 0.02 | 2015 | 44.119753 | -89.535683 |
| CPB2015-504C | WI | 0.09 | 2015 | 44.119753 | -89.535683 |
| CPBWGS_44 | WI (2022) | 5.02 | 2015 | 44.119753 | -89.535683 |
| CPBWGS_45 | WI (2022) | 4.92 | 2015 | 44.119753 | -89.535683 |
| CPBWGS_46 | WI (2022) | 3.61 | 2015 | 44.119753 | -89.535683 |
| CPBWGS_47 | WI (2022) | 4.63 | 2015 | 44.119753 | -89.535683 |
| CPBWGS_48 | WI (2022) | 5.28 | 2015 | 44.119753 | -89.535683 |

**Supplementary Table S2.** Average allele frequency per population between HiSeq and NovaSeq platforms.

| Population | HiSeq | NovaSeq | P-value, F (F crit), df (single factor ANOVA) |
|---|---|---|---|
| Long Island, NY | 0.228129 | 0.226375<br>0.226244<br>0.228486<br>0.225365 | 0.13, 4.2 (10.1), 4 |
| Hancock, WI | 0.229639 | 0.229678<br>0.230123<br>0.230014<br>0.234543 | 0.33, 1.3 (10.1), 4 |

**Supplementary Table S3.** Optimized parameter estimates for two-population divergence models. Models with constant population size ('no_mig') are favored over a model with fluctuating population size in descendent lineages ('no_mig_size').The optimized parameters (modern $N_e$ values and divergence time) are shown with and without mutation rate conversion.

| Model | Replicate | log-likelihood | AIC | AICc | $\chi^2$ | θ | Optimized params (*$N_{NYe}$, $N_{Wie}$, Divergence time in generations*, respectively*)* |
|---|---|---|---|---|---|---|---|
| no_mig | Round_4 | -43145.42 | 86296.84 | 86297.42 | 1012871.28 | 47493.23 | 2.3954, 5.7715, 0.0479 |

| | | | | | | | |
|---|---|---|---|---|---|---|---|
| | Replicate_38 | | | | | | 38660, 16045, 320 |
| no_mig | Round_4 Replicate_21 | -43146.19 | 86298.38 | 86298.96 | 1010975.13 | 47657.87 | 2.3175, 5.8832, 0.0471 |
| | | | | | | | 39545, 15577, 316 |
| no_mig | Round_4 Replicate_3 | -43147.57 | 86301.14 | 86301.73 | 821431.93 | 47513.32 | 2.1856, 6.0798, 0.0484 |
| | | | | | | | 40742, 14646, 324 |
| no_mig | Round_4 Replicate_34 | -43151.64 | 86309.28 | 86309.87 | 956893.33 | 47452.77 | 2.2246, 6.7254, 0.0482 |
| | | | | | | | 45011, 14888, 322 |
| no_mig | Round_4 Replicate_17 | -43152.07 | 86310.14 | 86310.73 | 927521.12 | 47624.83 | 2.1606, 6.6953, 0.0474 |
| | | | | | | | 44972, 14512, 318 |
| no_mig_size | Round_4 Replicate_22 | -46216.41 | 92444.82 | 92445.23 | 359258.73 | 49926.34 | 0.8645, 0.7619, 6.6653, 3.4216, 0.0102, 0.0275 |
| no_mig_size | Round_4 Replicate_2 | -46658.86 | 93329.72 | 93330.13 | 237048.27 | 50251.34 | 0.6412,1.0538,7.2372, 2.3909,0.0111,0.0268 |
| no_mig_size | Round_4 Replicate_12 | -46824.66 | 93661.32 | 93661.73 | 325138.65 | 50413.73 | 1.2038,0.9405,3.9139, 2.0381,0.0101,0.0265 |
| no_mig_size | Round_4 Replicate_30 | -47142.15 | 94296.30 | 94296.71 | 242894.59 | 48834.36 | 1.1876,0.6803,4.912, 2.8457,0.0104,0.0337 |
| no_mig_size | Round_4 Replicate_10 | -47402.98 | 94817.96 | 94818.37 | 298775.13 | 50844.37 | 0.7579,0.9602,9.5102, 1.8682,0.0101,0.0242 |

**Evidence of hard-selective sweeps suggest independent adaptation to insecticides in Colorado potato beetle (Coleoptera: *Chrysomelidae*) populations**

**Supplemental Table S4.** Genes in shared regions of high or low recombination (>10-fold difference in recombination rate relative to the background rate on each scaffold).

| **Genes in shared low recombination regions** | |
|---|---|
| **Gene** | **Annotation** |
| XP_023019051 | aquaporin AQPAe.a-like |
| XP_023012243 | uncharacterized protein LOC111502400 |
| XP_023025403 | uncharacterized protein F23F12.8-like |
| XP_023024874 | rho GTPase-activating protein 68F-like |
| XP_023020752 | zinc finger protein 888-like |
| XP_023021989 | equilibrative nucleoside transporter 3-like |
| XP_023019162 | uncharacterized protein LOC111507988 |
| XP_023019168<br>XP_023019167 | uncharacterized protein LOC111507993 isoform X1, X2 |
| XP_023018836 | venom carboxylesterase-6-like |
| XP_023017074 | zinc finger protein 142-like |
| XP_023012777 | flotillin-2 |
| XP_023012775 | glucose dehydrogenase |
| XP_023012782<br>XP_023012784 | dnaJ homolog subfamily C member 30-like |
| XP_023012785<br>XP_023012786 | serine/threonine-protein phosphatase PGAM5, mitochondrial isoform X1, X2 |
| XP_023022448 | alpha-tocopherol transfer protein-like |
| XP_023014614 | 60S ribosomal protein L22-like |
| XP_023018828 | DNA helicase MCM8-like |
| XP_023030429<br>XP_023030427<br>XP_023030428 | putative ammonium transporter 3 isoform X1, X2 |
| XP_023027679 | low molecular weight phosphotyrosine protein phosphatase 1-like |
| XP_023022952 | citron Rho-interacting kinase |
| XP_023029403<br>XP_023029404<br>XP_023029405<br>XP_023029406 | uncharacterized protein LOC111517468 isoform X1, X2 |
| XP_023016342 | putative sodium-coupled neutral amino acid transporter 11 |
| XP_023024439 | uncharacterized protein LOC111512529 |
| XP_023024291 | uncharacterized protein LOC111512404 |
| XP_023015429 | putative sphingolipid delta(4)-desaturase/C4-monooxygenase |
| XP_023012643<br>XP_023012644<br>XP_023012645 | putative tricarboxylate transport protein, mitochondrial isoform X1, X2 |
| XP_023013524 | protein phosphatase 1 regulatory subunit 15A |
| XP_023030020 | sperm-associated antigen 6-like |
| XP_023014145<br>XP_023014146<br>XP_023014147<br>XP_023014148 | tetratricopeptide repeat protein 14 isoform X1, X2, X3 |
| XP_023018118<br>XP_023018117<br>XP_023018119<br>XP_023018120 | zinc finger homeobox protein 3 isoform X1, X2, X3, X4 |
| XP_023021262<br>XP_023021261 | tafazzin homolog isoform X1, X2 |
| XP_023014784 | structure-specific endonuclease subunit slx1 |
| XP_023018027 | cilia- and flagella-associated protein 100-like |

| | |
|---|---|
| XP_023019070 | cytochrome P450 9e2-like |
| XP_023021162 | peroxidase-like |
| XP_023021163 | ubiquitin carboxyl-terminal hydrolase 7 |
| XP_023021164 | sulfate permease 2-like |
| XP_023014564<br>XP_023017889 | alanyl-tRNA editing protein Aarsd1 isoform X1, X2 |
| XP_023016356 | ficolin-2-like |
| XP_023029628 | titin |
| XP_023016616 | uncharacterized protein LOC111513564 |
| XP_023016615 | uncharacterized protein LOC111515999 |
| XP_023018236 | venom dipeptidyl peptidase 4-like |
| XP_023014427 | probable multidrug resistance-associated protein lethal(2)03659 |
| XP_023025556 | mitochondrial ornithine transporter 1 |
| XP_023027958 | monocarboxylate transporter 12-like |
| XP_023024728 | uncharacterized protein LOC111504309 |
| XP_023021441 | cathepsin L-like proteinase |
| XP_023028360 | uncharacterized protein LOC111502827 |
| XP_023017564 | peptidyl-prolyl cis-trans isomerase CWC27 homolog |
| XP_023014593 | uncharacterized protein LOC111509564 isoform X1 |
| XP_023012767 | multidrug resistance protein homolog 49-like |
| XP_023012759 | voltage-dependent calcium channel subunit alpha-2/delta-3-like |
| XP_023014594 | peroxisomal biogenesis factor 3 |
| XP_023021095 | gastrula zinc finger protein XlCGF71.1-like |
| XP_023021863 | oocyte zinc finger protein XlCOF19-like |
| XP_023027823 | probable 2-oxoglutarate dehydrogenase E1 component DHKTD1 homolog, mitochondrial |
| XP_023018198 | uncharacterized protein LOC111505511 |
| XP_023017170 | serine/threonine-protein kinase TBK1-like |
| XP_023026889 | tRNA (cytosine(34)-C(5))-methyltransferase |
| XP_023026274 | tetratricopeptide repeat protein 25 |
| XP_023016659 | venom protease-like |
| XP_023016104 | uncharacterized protein LOC111516144 |
| XP_023014530 | ganglioside-induced differentiation-associated protein 2-like |
| XP_023014531 | renin receptor |
| XP_023019078 | ATPase ASNA1 homolog |
| XP_023021787<br>XP_023028092 | receptor expression-enhancing protein 5-like |
| XP_023020346<br>XP_023020919<br>XP_023020918<br>XP_023023436<br>XP_023023438 | mitogen-activated protein kinase kinase kinase 4 isoform X1, X2, X3 |

| | |
|---|---|
| XP_023025193 | probable ATP-dependent RNA helicase DDX46 |
| XP_023025195 | uncharacterized protein LOC111506092 |
| XP_023025196 | calpain-7-like |
| XP_023025197<br>XP_023025198 | uncharacterized protein LOC111505839 isoform X1, X2 |
| XP_023022065 | vesicle transport through interaction with t-SNAREs homolog 1B |
| XP_023016804 | RNA-binding protein pno1 |
| XP_023013674 | protein phosphatase 1 regulatory subunit 21 |
| XP_023016484 | uncharacterized protein LOC111506887 |
| XP_023016483 | GPN-loop GTPase 3 |
| XP_023019509 | uncharacterized protein LOC111510793 |
| XP_023019497 | Niemann-Pick type protein homolog 1B-like |
| XP_023019485 | uncharacterized protein LOC111503147 |
| XP_023017876<br>XP_023029640 | venom protease-like isoform X1, X2 |
| XP_023022506 | ankyrin repeat, SAM and basic leucine zipper domain-containing protein 1-like |
| XP_023012615 | xanthine dehydrogenase-like |
| XP_023013142 | JNK1/MAPK8-associated membrane protein |
| XP_023025017 | serine/threonine-protein phosphatase 2A 56 kDa regulatory subunit epsilon isoform |
| XP_023025016 | early growth response protein 1-B |
| XP_023025011 | proliferation-associated protein 2G4 |
| XP_023019888 | histone acetyltransferase KAT7 |
| XP_023022504 | protein penguin |
| XP_023022501 | glycine cleavage system H protein, mitochondrial |
| XP_023025897 | gamma-aminobutyric acid receptor subunit beta |
| XP_023018499 | glutamate receptor 1 |
| XP_023016387 | ras-related and estrogen-regulated growth inhibitor |
| XP_023026299<br>XP_023012191 | uncharacterized protein LOC111502317 isoform X1, X2 |
| XP_023012203 | glutamate receptor ionotropic, NMDA 2D-like |
| XP_023012213<br>XP_023012226<br>XP_023012151 | probable cationic amino acid transporter isoform X1, X2, X3 |
| XP_023012159 | protein furry |
| XP_023012139<br>XP_023022063<br>XP_023022062<br>XP_023022061 | ubiquitin carboxyl-terminal hydrolase 35 isoform X1, X2 |
| XP_023011562<br>XP_023027251<br>XP_023027233<br>XP_023027238<br>XP_023027245 | neuronal PAS domain-containing protein 2-like isoform X1, X2, X3, X4 |

| | |
|---|---|
| XP_023027357<br>XP_023027363 | transmembrane protein 192 isoform X1, X2 |
| XP_023027350<br>XP_023027371 | tubulin polyglutamylase TTLL4 isoform X1, X2 |
| XP_023027378 | N-acetylneuraminate lyase-like |
| XP_023027597 | probable Dol-P-Man:Man(7)GlcNAc(2)-PP-Dol alpha-1,6-mannosyltransferase |
| XP_023027604<br>XP_023027613<br>XP_023027620<br>XP_023027629<br>XP_023027637 | high affinity cAMP-specific and IBMX-insensitive 3',5'-cyclic phosphodiesterase 8 isoform X1, X2, X3, X4, X5 |
| XP_023023605 | short coiled-coil protein B-like isoform X2 |
| XP_023023620 | short coiled-coil protein homolog isoform X1 |
| XP_023023612 | 26S proteasome regulatory subunit 4 |
| XP_023023599 | vicilin-like seed storage protein At2g18540 |
| XP_023023626 | uncharacterized protein LOC111506981 |
| XP_023023669 | zinc finger protein 271-like isoform X2 |
| XP_023023661 | zinc finger protein 829-like isoform X1 |
| XP_023023677 | probable ATP-dependent RNA helicase Dbp73D |
| XP_023030505 | STE20-related kinase adapter protein alpha |
| XP_023017970 | zinc transporter 2-like isoform X1 |
| XP_023025948 | LIM and SH3 domain protein Lasp |
| XP_023025946 | 3-hydroxyisobutyrate dehydrogenase, mitochondrial |
| XP_023025923 | probable myosin heavy chain ECU04 1000 |
| XP_023025936 | MAP/microtubule affinity-regulating kinase 4-like |
| XP_023016514<br>XP_023016511 | integrin alpha-PS4-like isoform X1, X2 |
| XP_023016510<br>XP_023017959 | uncharacterized protein LOC111504253 isoform X1, X2 |
| XP_023015174 | translation elongation factor 2 |
| XP_023030093<br>XP_023030094 | protein BTG3-like |
| XP_023014539 | uncharacterized protein LOC111518071 |
| XP_023014544 | transcription factor BTF3 homolog 4 |
| XP_023014461<br>XP_023014444 | translocator protein-like |
| XP_023014452 | uncharacterized protein LOC111517287 |
| XP_023030180 | dopamine receptor 1 |
| XP_023017792<br>XP_023017791 | nuclear receptor coactivator 6-like isoform X1, X2 |

### Evidence of hard-selective sweeps suggest independent adaptation to insecticides in Colorado potato beetle (Coleoptera: *Chrysomelidae*) populations

| | |
|---|---|
| XP_023017790<br>XP_023029147<br>XP_023023943<br>XP_023017334 | equilibrative nucleoside transporter 3-like isoform X1, X2 |
| XP_023017336 | uncharacterized protein LOC111510559 isoform X3 |
| XP_023022249 | probable serine incorporator |
| XP_023022248 | uncharacterized protein LOC111502322 |
| XP_023022245 | uncharacterized protein LOC111515591 |
| XP_023022246 | uncharacterized protein LOC111516064 |
| XP_023022244 | probable histone-lysine N-methyltransferase PRDM7 |
| XP_023027073 | uncharacterized protein LOC111516178 |
| XP_023012156<br>XP_023027573<br>XP_023028017<br>XP_023028715 | uncharacterized protein LOC111504252 isoform X1, X2, X3, X4 |
| XP_023028123 | putative fatty acyl-CoA reductase CG5065 |
| XP_023014535 | uncharacterized protein LOC111506198 |
| XP_023014537 | multidrug resistance-associated protein 4-like |
| XP_023014536 | potassium voltage-gated channel protein eag |
| XP_023014534 | programmed cell death protein 6 |
| XP_023015361 | Bardet-Biedl syndrome 7 protein homolog |
| XP_023016973 | 39S ribosomal protein L23, mitochondrial |
| XP_023011971 | uncharacterized protein LOC111509119 |
| XP_023011972 | protein ZBED8-like |
| XP_023022884 | serine, glycine and glutamine-rich protein-like |
| XP_023019161 | activating transcription factor 3 |
| XP_023018079 | ubiquitin-like protein 3 |
| XP_023018081<br>XP_023020581 | transcription factor SOX-13 |
| XP_023018416 | fatty acid synthase-like |
| XP_023015796 | pre-mRNA cleavage complex 2 protein Pcf11-like |
| XP_023014063<br>XP_023021330 | leucine-rich repeat transmembrane neuronal protein 4-like |
| XP_023030420<br>XP_023030421<br>XP_023029483 | copper-transporting ATPase 1 isoform X1, X2, X3 |
| XP_023024528 | phosphatidylinositol 5-phosphate 4-kinase type-2 alpha-like |
| XP_023018429<br>XP_023018436<br>XP_023018681 | WD repeat-containing protein 55 homolog |
| XP_023018689 | AP-1 complex subunit beta-1-like |
| XP_023018697 | DNA repair protein REV1 |
| XP_023023570 | uncharacterized protein LOC111509609 |
| XP_023026734 | tyrosine-protein phosphatase non-receptor type 11-like |

| | |
|---|---|
| XP_023028529 | glutathione hydrolase 1 proenzyme-like isoform X2 |
| XP_023028901 | PAN2-PAN3 deadenylation complex catalytic subunit PAN2-like |
| XP_023026111 | peptidyl-prolyl cis-trans isomerase 5 |
| XP_023016559 | uncharacterized protein LOC111516249 |
| XP_023021148 | uncharacterized protein LOC111515504 |
| XP_023021149<br>XP_023020657<br>XP_023021811<br>XP_023015039<br>XP_023028195<br>XP_023027496<br>XP_023015422<br>XP_023015423<br>XP_023015425 | inter-alpha-trypsin inhibitor heavy chain H4-like isoform X1, X2, X3, X4, X5, X6, X7, X8 |
| XP_023015426 | vacuolar protein sorting-associated protein 13-like |
| XP_023015424 | protein doublesex-like |
| XP_023015430 | uncharacterized protein LOC111515195 |
| XP_023015427 | DNA replication licensing factor Mcm7 |
| XP_023015428 | 60S ribosomal protein L8 |
| XP_023015431<br>XP_023022780 | putative U5 small nuclear ribonucleoprotein 200 kDa helicase |
| XP_023024433 | muscleblind-like protein 1 |
| XP_023027196 | multidrug resistance-associated protein 4-like |
| XP_023027703 | ficolin-2-like |
| XP_023026218<br>XP_023028749<br>XP_023027359 | uncharacterized protein LOC111517784 isoform X1, X2, X3 |
| XP_023024222 | glutaredoxin-related protein 5, mitochondrial |
| XP_023027692 | prostatic acid phosphatase-like |
| XP_023023915<br>XP_023029829 | uncharacterized protein LOC111503810 isoform X1, X2 |
| XP_023029828<br>XP_023029827<br>XP_023029821 | insulin-degrading enzyme isoform X1, X2, X3 |
| XP_023016140 | lachesin |
| XP_023013988 | solute carrier family 22 member 4-like |
| XP_023013987<br>XP_023021589<br>XP_023021581 | formin-binding protein 1-like isoform X1, X2, X3 |
| XP_023021573 | protein disulfide-isomerase A5 |
| XP_023012260 | delta-1-pyrroline-5-carboxylate synthase |
| XP_023026324 | splicing factor 1 |
| XP_023013125<br>XP_023013124 | venom serine carboxypeptidase-like |
| XP_023013123 | multidrug resistance-associated protein 4-like isoform X2 |
| XP_023019076 | probable multidrug resistance-associated protein lethal(2)03659 isoform X1 |

| | |
|---|---|
| XP_023021504<br>XP_023022381 | T-cell activation inhibitor, mitochondrial isoform X1, X2 |
| XP_023012514 | uncharacterized protein LOC111505964 |
| XP_023012513<br>XP_023024499 | polygalacturonase-like isoform X1, X2 |
| XP_023024491 | CWF19-like protein 2 homolog |
| XP_023025345 | stromal interaction molecule homolog |
| XP_023025344 | RING finger protein 121 |
| XP_023016650 | probable cytochrome P450 6a23 |
| XP_023016649 | LIM domain transcription factor LMO4.2-like |
| XP_023016648 | uncharacterized protein LOC111503173 |
| XP_023015868 | uncharacterized protein LOC111518025 |
| XP_023015818<br>XP_023029083 | uncharacterized protein LOC111505501 isoform X1, X2 |
| XP_023024022 | probable multidrug resistance-associated protein lethal(2)03659 isoform X2 |
| XP_023022391 | uncharacterized protein LOC111515499 |
| XP_023022392 | uncharacterized protein LOC111515125 |
| XP_023013186 | uncharacterized protein LOC111512767 |
| XP_023013177 | uncharacterized protein LOC111512766 |
| XP_023030113 | uncharacterized protein LOC111510492 |
| XP_023016095<br>XP_023016094 | beta-catenin-like protein 1 |
| XP_023023852 | multidrug resistance-associated protein 4-like |
| XP_023027492 | soluble guanylate cyclase 88E |
| XP_023027128 | solute carrier family 22 member 4-like |
| XP_023024688<br>XP_023024687<br>XP_023022185 | adrenodoxin-like protein, mitochondrial |
| XP_023020796 | lamin-B receptor |
| XP_023020797 | heterogeneous nuclear ribonucleoprotein 87F |
| XP_023012969<br>XP_023029751 | B-cell lymphoma/leukemia 11A isoform X1, X2 |
| XP_023028064<br>XP_023020729<br>XP_023020730<br>XP_023020731<br>XP_023022099 | disks large 1 tumor suppressor protein isoform X1, X2, X3, X4, X5 |
| XP_023026677<br>XP_023018745<br>XP_023018746<br>XP_023022564 | uncharacterized protein LOC111503036 isoform X1, X2, X3, X4 |
| XP_023022356 | uncharacterized protein LOC111508564 |

| | |
|---|---|
| XP_023022488 | cell division control protein 42 homolog |
| XP_023022426 | corepressor interacting with RBPJ 1-like |
| XP_023022283 | uncharacterized protein LOC111510097 |
| XP_023013003 | UPF0545 protein C22orf39 homolog |
| XP_023013002 | intraflagellar transport protein 52 homolog |
| XP_023013005 | protein phosphatase 1E-like |
| XP_023013004 | uncharacterized MFS-type transporter C09D4.1-like |
| XP_023019881 | protein OPI10 homolog |
| XP_023018807 | notchless protein homolog 1 |
| XP_023022358 | venom allergen 3-like isoform X4 |
| XP_023021725 | putative ATPase N2B |
| XP_023030129 | prefoldin subunit 3 |
| XP_023030127 | feline leukemia virus subgroup C receptor-related protein 2-like |
| XP_023030126 | lysine-specific histone demethylase 1A-like |
| XP_023024267<br>XP_023015801 | dynein beta chain, ciliary-like |
| XP_023015802 | uncharacterized protein LOC111508923 |
| XP_023015806 | potassium voltage-gated channel subfamily H member 8-like |
| XP_023015809 | uncharacterized protein LOC111501979 |
| XP_023019587 | syntenin-1-like |
| XP_023024847 | mediator of RNA polymerase II transcription subunit 19 |
| XP_023017151 | NADPH--cytochrome P450 reductase |
| XP_023024904 | odorant receptor 4-like |
| XP_023023241<br>XP_023020334<br>XP_023024156 | regulator of chromosome condensation isoform X1, X2 |
| XP_023011683 | eukaryotic translation initiation factor 6 |
| XP_023018486<br>XP_023018483<br>XP_023016130 | serine/threonine-protein kinase MARK2-like isoform X12 |
| XP_023019277 | laminin subunit gamma-1-like |
| XP_023017406<br>XP_023017407 | rutC family protein UK114 isoform X2, X4 |
| XP_023017408 | cell adhesion molecule 3-like |
| XP_023017405 | cytochrome P450 6a2-like |
| XP_023029713<br>XP_023029714<br>XP_023029715<br>XP_023013323<br>XP_023021542 | proton-coupled amino acid transporter 4-like isoform X1, X2, X3, X4 |
| XP_023021540 | structural maintenance of chromosomes protein 1A |
| XP_023013061<br>XP_023020273 | F-actin-uncapping protein LRRC16A isoform X1, X2 |
| XP_023020274 | adenylate kinase isoenzyme 6 |
| XP_023018377 | uncharacterized protein LOC111502544 |

| | |
|---|---|
| XP_023018378 | uncharacterized protein LOC111513637 |
| XP_023018379 | C-type lectin mannose-binding isoform-like |
| XP_023018381 | perlucin-like protein |
| XP_023018382 | myosin light chain kinase, smooth muscle-like |
| XP_023022814 | uncharacterized protein LOC111503801 |
| XP_023024105 | la-related protein 6 |
| XP_023024113<br>XP_023024145<br>XP_023012413 | polyadenylate-binding protein 2 isoform X1, X2, X3 |
| XP_023025627 | uncharacterized protein LOC111513684 |
| XP_023015124<br>XP_023025820 | facilitated trehalose transporter Tret1-like isoform X1, X2 |
| XP_023019109 | uncharacterized protein LOC111509486 |
| XP_023013982 | uncharacterized protein PF11_0213-like |
| XP_023030494 | uncharacterized protein LOC111510351 |
| XP_023019732 | nucleolar protein 58 |
| XP_023019729 | peroxisome proliferator-activated receptor gamma coactivator 1-alpha |
| XP_023019731 | peroxisome proliferator-activated receptor gamma coactivator 1-alpha |
| XP_023025679 | ABC transporter G family member 20 |
| XP_023024767 | ABC transporter G family member 20 |
| XP_023024768 | ABC transporter G family member 20 |
| XP_023021013 | uncharacterized protein LOC111508382 |
| XP_023021008 | uncharacterized protein LOC111515059 |
| XP_023022023 | uncharacterized protein LOC111516846 |
| XP_023022866 | uncharacterized protein LOC111507428 |
| XP_023024188 | uncharacterized protein LOC111507428 |
| XP_023024189 | uncharacterized protein LOC111516461 |
| XP_023014771 | uncharacterized protein LOC111515564 |
| XP_023014780 | neprilysin-4-like |
| XP_023014788 | nose resistant to fluoxetine protein 6-like |
| XP_023019618 | uncharacterized protein C7orf50 homolog |
| XP_023027063 | putative aminopeptidase W07G4.4 |
| XP_023028714 | cGMP-specific 3',5'-cyclic phosphodiesterase-like |
| LDEC001798-RA | myb sant-like transcription factor |
| LDEC001799-RA | nuclease harbi1 |
| LDEC002010-RA | tigger transposable element-derived protein 1-like isoform x1\|GO:0003677 |
| LDEC002042-RA | PREDICTED: uncharacterized protein LOC105460119, partial |

| | |
|---|---|
| LDEC003497-RA<br>LDEC003497-RB | dnaj homolog subfamily c member 30 |
| LDEC003953-RA | inosine-uridine preferring nucleoside hydrolase-like protein |
| LDEC005290-RA | hypothetical protein X777_07635\|GO:0005488 |
| LDEC005740-RA | reverse ribonuclease integrase\|GO:0005488 GO:0043170 GO:0044238 |
| LDEC006598-RA | PREDICTED: uncharacterized protein LOC105556339, partial |
| LDEC007886-RA | 115 kDa in type-1 retrotransposable element R1DM |
| LDEC008137-RA | cAMP-dependent kinase type I regulatory subunit isoform X1\|GO:0005952 GO:0008603 GO:0016301 GO:0045859 |
| LDEC008320-RA | zinc finger dna binding protein |
| LDEC009403-RA | zinc-finger associated domain containing partial\|GO:0005634 GO:0008270 |
| LDEC010108-RA | reverse ribonuclease partial |
| LDEC010109-RA | reverse ribonuclease partial\|GO:0003676 GO:0015074 |
| LDEC013334-RA | cytochrome b-c1 complex subunit 8 |
| LDEC014205-RA | rna-directed dna polymerase from mobile element partial |
| LDEC015217-RA | hypothetical protein ALC57_01510 |
| LDEC015249-RA | transcription factor adf-1-like isoform x2 |
| LDEC015498-RA | 60s ribosomal protein l8\|GO:0015934 GO:0003723 GO:0003735 GO:0006412 GO:0042254 |
| LDEC015653-RA | retrovirus-related pol polyprotein from transposon tnt 1-94 |
| LDEC015654-RA | hypothetical protein RP20_CCG014482 |
| LDEC015687-RA | retrovirus-related pol polyprotein from transposon tnt 1-94 |
| LDEC015896-RA | piggybac transposable element-derived protein 4-like |
| LDEC016262-RA | nuclease harbi1 |
| LDEC016458-RA | inactive dipeptidyl peptidase 10\|GO:0016021 GO:0008236 GO:0006508 |
| LDEC016587-RA | potassium voltage-gated channel subfamily h member 8 isoform x6\|GO:0006811 GO:0044763 |
| LDEC017187-RA | retrovirus-related pol polyprotein from transposon partial |
| LDEC017728-RA | acetylcholine receptor subunit alpha-like isoform x2\|GO:0016021 GO:0030054 GO:0045211 GO:0004889 GO:0098655 GO:0007165 |
| LDEC018288-RA | hypothetical protein EAG_04425, partial |
| LDEC018355-RA | rna-directed dna polymerase from mobile element jockey-like |
| LDEC018459-RA | nuclease harbi1 |
| LDEC018460-RA | nuclease harbi1 |
| LDEC018561-RA | Zinc finger 862 |
| LDEC019168-RA | PREDICTED: uncharacterized protein LOC107218977, partial |
| LDEC020862-RA | PO21_NASVIRecName: Full=Retrovirus-related Pol polyprotein from type-1 retrotransposable element R2; AltName: Full=Retrovirus-related Pol polyprotein from type I retrotransposable element R2; Includes: RecName: Full=Reverse transcriptase; Includes: RecName: Full=Endonuclease |
| LDEC021439-RA | reverse partial\|GO:0003676 GO:0008270 |
| LDEC021475-RA | gustatory receptor |
| LDEC021647-RA | zinc finger MYM-type 2 isoform X1 |
| LDEC021774-RA | hypothetical protein D910_02092\|GO:0005488 |

| | |
|---|---|
| LDEC022096-RA | PREDICTED: uncharacterized protein LOC105663446 |
| LDEC022133-RA | hypothetical protein YQE_12498, partial |
| LDEC022414-RA | zinc finger protein 862\|GO:0016020 |
| LDEC022838-RA | piggybac transposable element-derived protein 4-like |
| LDEC022958-RA | hypothetical protein TcasGA2_TC012951 |
| LDEC023393-RA | reverse ribonuclease partial\|GO:0003676 GO:0015074 |
| LDEC023730-RA | gag-pol polyprotein |
| LDEC023731-RA | PREDICTED: uncharacterized protein LOC105195918 |
| LDEC024167-RA | wd repeat-containing protein 75 |
| LDEC024217-RA | PREDICTED: uncharacterized protein LOC103314333 |
| LDEC024383-RA | ATPase N2B\|GO:0005524 |
| **Genes in shared high recombination regions** | |
| XP_023026553 | venom carboxylesterase-6-like |
| XP_023030196 | type I inositol 3,4-bisphosphate 4-phosphatase |
| XP_023019473 | J domain-containing protein |

**Evidence of hard-selective sweeps suggest independent adaptation to insecticides in Colorado potato beetle (Coleoptera: *Chrysomelidae*) populations**

**Supplementary Table S5**. Genes identified as candidates of hard selective sweeps, asterisk denotes genes also significant in Pélissié et al., 2021 from PCAdapt.

| **Gene ID** | **Protein Name** |
|---|---|
| **Long Island, New York** | |
| XP_023012173, XP_023012180, LDEC004451 | acetyl-coenzyme A synthetase isoform X1, X2 |
| XP_023014225 | paired mesoderm homeobox protein 2A-like |
| XP_023018886 | E3 ubiquitin-protein ligase HECW2 isoform X1 |
| XP_023018894 | E3 ubiquitin-protein ligase HECW2 isoform X2 |
| XP_023022462 | tctex1 domain-containing protein 2-like |
| XP_023023146<br>XP_023023153<br>XP_023023162<br>XP_023023173<br>XP_023023180 | WD repeat-containing protein 47 isoform X1, X2, X4, X5 |
| XP_023024291 | uncharacterized protein LOC111512404 |
| XP_023029010 | G protein-coupled receptor kinase 2 |
| XP_023029080 | glutamate receptor ionotropic, kainate 2-like |
| XP_023029495 | uncharacterized protein LOC111517537 |
| XP_023012335<br>XP_023012336 | sarcosine dehydrogenase, mitochondrial isoform X1, X2 |
| XP_023012612 | phytanoyl-CoA dioxygenase, peroxisomal-like |
| XP_023012615 | Niemann-Pick type protein homolog 1B-like |
| XP_023013000 | hrp65 protein-like |
| XP_023013577 | uncharacterized protein LOC111503500 |
| *XP_023013985 & LDEC018602 | neuronal acetylcholine receptor subunit alpha-7-like |
| XP_023017816<br>XP_023017817<br>XP_023017818 | cerebellar degeneration-related protein 2 isoform X1, X2, X3 |
| XP_023018113<br>XP_023018114 | cytochrome P450 9e2-like isoform X1, X2 |
| XP_023018115 | cytochrome P450 9e2-like |

| | |
|---|---|
| XP_023018337 | zinc finger protein rotund-like |
| XP_023018340 | putative DNA helicase Ino80 |
| XP_023018574<br>XP_023018575 | uncharacterized protein LOC111507501 isoform X1, X2 |
| XP_023018576<br>XP_023018577<br>XP_023018578 | receptor expression-enhancing protein 1 isoform X2, X3, X4 |
| XP_023019275 | small conductance calcium-activated potassium channel protein |
| XP_023019639 | retinaldehyde-binding protein 1-like isoform X1 |
| XP_023019641 | alpha-tocopherol transfer protein-like isoform X2 |
| XP_023019711 | poly(A) polymerase type 3-like |
| XP_023020189<br>XP_023020190<br>XP_023020191<br>XP_023020192<br>XP_023020193 | palmitoyltransferase ZDHHC15 isoform X1, X2, X3, X4, X5 |
| XP_023020349 | uncharacterized protein LOC111508933 |
| XP_023020650 | mucin-5AC-like |
| XP_023020747 | esterase FE4-like |
| XP_023020828 | glycine receptor subunit alpha-2 |
| XP_023022114<br>XP_023022115<br>XP_023022116 | MOXD1 homolog 1 isoform X1, X2 |
| XP_023022146 | equilibrative nucleoside transporter 3-like |
| XP_023022258 | homeobox protein onecut |
| XP_023022877 | protein phosphatase 1 regulatory subunit 12B-like |
| XP_023023108<br>XP_023023109<br>XP_023023110<br>XP_023023111 | deoxynucleoside kinase-like isoform X1, X2 (x3) |
| XP_023023329 | protein ALP1-like |

| | |
|---|---|
| XP_023023651 | uncharacterized protein LOC111511867 |
| XP_023024118 | arfGAP with SH3 domain, ANK repeat and PH domain-containing protein |
| XP_023024473 | sensory neuron membrane protein 2-like |
| XP_023024697 | mitochondrial amidoxime-reducing component 1-like |
| XP_023025254 | uncharacterized protein LOC111513291 |
| XP_023025815 | neuroendocrine convertase 2-like |
| XP_023026000 | uncharacterized protein LOC111514008 |
| XP_023026103 | uncharacterized protein LOC111514100 |
| XP_023026304 | short stature homeobox protein 2-like |
| XP_023026446 | carboxypeptidase Q-like |
| XP_023026654 | uncharacterized protein LOC111514631 isoform X2 |
| XP_023026681 | protein ABHD17B |
| XP_023026978 | multidrug resistance-associated protein 1-like |
| XP_023027200 | dnaJ protein homolog 1-like |
| XP_023027618 | GMP synthase [glutamine-hydrolyzing] |
| XP_023028679 | uncharacterized protein LOC111516799 |
| *LDEC000495 | uncharacterized protein |
| LDEC001211 | uncharacterized protein |
| *LDEC002091 | dna-binding protein p3a2 isoform x1 |
| LDEC002092 | uncharacterized protein |
| LDEC003243 | PREDICTED: uncharacterized protein LOC106134883 |
| LDEC003725 | transposable element p transposase |
| *LDEC003903 | 52 kda repressor of the inhibitor of the protein kinase- partial |
| *LDEC003933 | hypothetical protein ALC57_16739 |

| | |
|---|---|
| LDEC005595 | PREDICTED: uncharacterized protein LOC105379985 |
| LDEC007235 | uncharacterized protein |
| *LDEC007352 | Zinc finger rotund |
| LDEC007841 | Eukaryotic translation initiation factor 4 gamma 1 |
| LDEC008162 | ATP-dependent DNA helicase PIF1-like |
| *LDEC009455 | tigger transposable element-derived protein 6-like |
| *LDEC010707 | rna-directed dna polymerase from mobile element jockey-like |
| LDEC010826 | PREDICTED: uncharacterized protein LOC105570862 |
| LDEC011938 | arf-gap with sh3 ank repeat and ph domain-containing protein 2 isoform x1 |
| LDEC012523 | moxd1 homolog 1 |
| LDEC013064 | set and mynd domain-containing protein 4 |
| LDEC014604 | partitioning defective 3 homolog isoform x5 |
| LDEC014808 | PREDICTED: uncharacterized protein LOC103519749 |
| *LDEC015662 | alpha beta hydrolase domain-containing protein 17b |
| LDEC016056 | phytanoyl- peroxisomal-like |
| *LDEC016065 | niemann-pick c1 protein isoform x1 |
| *LDEC016157 | hypothetical protein TcasGA2_TC001793 |
| *LDEC016177 | one cut domain family member 2 |
| *LDEC016194 | paired box protein |
| LDEC017263 | deoxynucleoside kinase-like |
| *LDEC017782 | lachesin isoform x2 |
| *LDEC018458 | myb sant-like transcription factor |
| LDEC018839 | 115 kDa in type-1 retrotransposable element R1DM |
| *LDEC019773 | uncharacterized protein |
| LDEC021148 | uncharacterized protein |
| LDEC021495 | PREDICTED: uncharacterized protein LOC107265388 |

| | |
|---|---|
| *LDEC022445 | retrotransposable element tf2 155 kda protein type 1 |
| LDEC024532 | uncharacterized protein |
| **Hancock, Wisconsin** | |
| XP_023029844 | uncharacterized protein LOC111517795 |
| XP_023014225 | paired mesoderm homeobox protein 2A-like |
| XP_023017838 | latrophilin Cirl |
| XP_023018886<br>XP_023018894 | E3 ubiquitin-protein ligase HECW2 isoform X1, X2 |
| XP_023025569 | rho GTPase-activating protein 190 |
| XP_023026588 | zinc finger SWIM domain-containing protein 8-like |
| XP_023026644<br>XP_023026650 | heparan sulfate glucosamine 3-O-sulfotransferase 1 |
| XP_023026945 | homeotic protein proboscipedia |
| XP_023028564 | zinc finger protein 271-like |
| XP_023029795 | uncharacterized protein LOC111517764 |
| XP_023030025 | protein 5NUC-like |
| *XP_023011673<br>LDEC007110 | ATP-binding cassette sub-family D member 3 |
| XP_023011806 | lachesin-like |
| XP_023012143 | histone deacetylase 7 |
| XP_023012615 | Niemann-Pick type protein homolog 1B-like |
| XP_023012643<br>XP_023012644<br>XP_023012645 | putative tricarboxylate transport protein, mitochondrial isoform X1,X2 |

| | |
|---|---|
| XP_023013101 | semaphorin-1A |
| XP_023016418<br>XP_023016419 | alpha-crystallin B chain-like isoform X1, X2 |
| XP_023016920 | segmentation protein cap'n'collar |
| XP_023017638<br>XP_023017639<br>XP_023017640, XP_023017641,<br>XP_023017642 | glutaredoxin domain-containing cysteine-rich protein CG31559-like isoform X1, X2 |
| XP_023018172 | uncharacterized protein LOC111507143 |
| XP_023019181 | zinc finger protein 148-like |
| XP_023020349 | uncharacterized protein LOC111508933 |
| XP_023020608 | uncharacterized protein LOC111509151 |
| XP_023022354 | gustatory and odorant receptor 24 |
| XP_023022657 | putative nuclease HARBI1 |
| XP_023022775 | organic cation transporter protein-like |
| XP_023022805 | uncharacterized protein LOC111511026 |
| XP_023022884 | potassium voltage-gated channel protein eag |
| XP_023023418 | E3 ubiquitin-protein ligase SHPRH |
| XP_023023971 | chorion peroxidase |
| XP_023024026 | insulin gene enhancer protein ISL-1 |
| XP_023024383 | uncharacterized protein LOC111512483 |
| XP_023024611 | NGFI-A-binding protein homolog isoform X1 |
| XP_023024651 | TGF-beta-activated kinase 1 and MAP3K7-binding protein 2 |
| XP_023024750 | uncharacterized protein LOC111512816 |
| XP_023025254 | uncharacterized protein LOC111513291 |
| XP_023025360 | cathepsin B-like |
| XP_023025369 | uncharacterized protein LOC111513398 |

| | |
|---|---|
| XP_023025560 | uncharacterized protein LOC111513568 |
| XP_023025815 | neuroendocrine convertase 2-like |
| XP_023026421 | F-BAR domain only protein 2-like |
| XP_023026791 | uncharacterized protein LOC111514777 |
| XP_023026792 | LOW QUALITY PROTEIN: glucose dehydrogenase [FAD, quinone]-like |
| XP_023026978 | multidrug resistance-associated protein 1-like |
| XP_023027618 | LOW QUALITY PROTEIN: GMP synthase [glutamine-hydrolyzing] |
| XP_023027778 | uncharacterized protein LOC111515808 |
| *LDEC001490 | Insulin gene enhancer protein ISL-1 |
| *LDEC002030 | PREDICTED: uncharacterized protein LOC107883157 |
| *LDEC002578 | tricarboxylate transport mitochondrial |
| *LDEC003667 | chorion peroxidase isoform x1 |
| LDEC003718 | PREDICTED: uncharacterized protein LOC107883157 |
| *LDEC003881 | heparan sulfate glucosamine 3-o-sulfotransferase 1 isoform x1 |
| LDEC004568 | piggybac transposable element-derived protein 3-like |
| *LDEC005491 | ubiquinone biosynthesis monooxygenase coq6 |
| *LDEC006880 | bel12 ag transposon poly |
| LDEC009247 | hypothetical protein ALC62_00085, partial |
| *LDEC009411 | hydrocephalus-inducing protein |
| LDEC009412 | hydrocephalus-inducing protein homolog |
| LDEC011319 | 40s ribosomal protein s3a |
| LDEC011494 | glucose dehydrogenase |

| | |
|---|---|
| LDEC013739 | gustatory receptor 63a |
| LDEC015368 | transposable element tc3 partial |
| LDEC015768 | PREDICTED: uncharacterized protein LOC107883157 |
| LDEC016025 | hypothetical protein ALC62_00085, partial |
| *LDEC016065 | niemann-pick c1 protein isoform x1 |
| *LDEC016194 | paired box protein |
| LDEC016943 | salicyl alcohol oxidase |
| *LDEC017124 | organic cation transporter protein |
| LDEC017703 | uncharacterized protein |
| LDEC020790 | uncharacterized protein |
| LDEC021495 | PREDICTED: uncharacterized protein LOC107265388 |
| *LDEC022531 | esterase |

**Evidence of hard-selective sweeps suggest independent adaptation to insecticides in Colorado potato beetle (Coleoptera: *Chrysomelidae*) populations**

**Supplemental Table S6.** GO terms for shared significantly swept loci (Hancock, WI and Long Island, NY)

| | go_id | go_name | root_node |
|---|---|---|---|
| 1 | GO:0005975 | carbohydrate metabolic process | biological_process |
| 2 | GO:0005694 | chromosome | cellular_component |
| 3 | GO:0004252 | serine-type endopeptidase activity | molecular_function |
| 4 | GO:0005634 | nucleus | cellular_component |
| 5 | GO:0005615 | extracellular space | cellular_component |
| 6 | GO:0007155 | cell adhesion | biological_process |
| 7 | GO:0008234 | cysteine-type peptidase activity | molecular_function |
| 8 | GO:0006177 | GMP biosynthetic process | biological_process |
| 9 | GO:0016021 | integral component of membrane | cellular_component |
| 10 | GO:0003964 | RNA-directed DNA polymerase activity | molecular_function |
| 11 | GO:0003922 | GMP synthase (glutamine-hydrolyzing) activity | molecular_function |
| 12 | GO:0005484 | SNAP receptor activity | molecular_function |
| 13 | GO:0006820 | anion transport | biological_process |
| 14 | GO:0016462 | pyrophosphatase activity | molecular_function |
| 15 | GO:0071260 | cellular response to mechanical stimulus | biological_process |
| 16 | GO:0016192 | vesicle-mediated transport | biological_process |
| 17 | GO:0055085 | transmembrane transport | biological_process |
| 18 | GO:0006541 | glutamine metabolic process | biological_process |
| 19 | GO:0046872 | metal ion binding | molecular_function |
| 20 | GO:0035556 | intracellular signal transduction | biological_process |
| 21 | GO:0006508 | proteolysis | biological_process |
| 22 | GO:0006278 | RNA-dependent DNA biosynthetic process | biological_process |

### Evidence of hard-selective sweeps suggest independent adaptation to insecticides in Colorado potato beetle (Coleoptera: *Chrysomelidae*) populations

| | | | |
|---|---|---|---|
| 23 | GO:0061025 | membrane fusion | biological_process |
| 24 | GO:0050790 | regulation of catalytic activity | biological_process |
| 25 | GO:0005829 | cytosol | cellular_component |
| 26 | GO:0005525 | GTP binding | molecular_function |
| 27 | GO:0005509 | calcium ion binding | molecular_function |

**Evidence of hard-selective sweeps suggest independent adaptation to insecticides in Colorado potato beetle (Coleoptera: *Chrysomelidae*) populations**

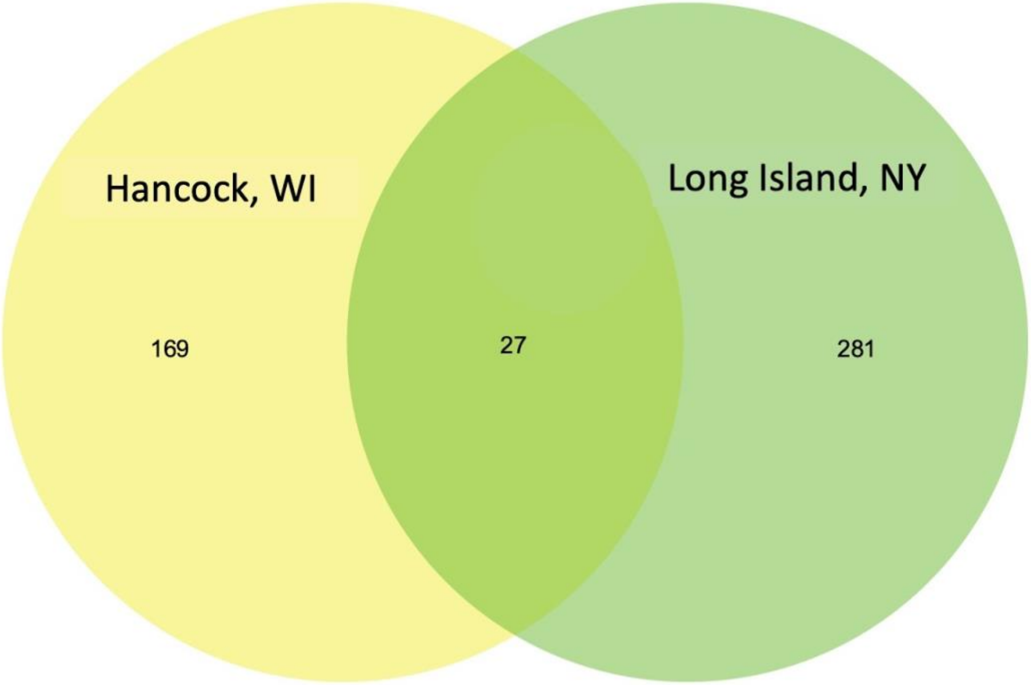


**Supplemental Figure S1.** Venn diagram showing overlap in enriched GO terms for HAN and LI.

**Evidence of hard-selective sweeps suggest independent adaptation to insecticides in Colorado potato beetle (Coleoptera: *Chrysomelidae*) populations**

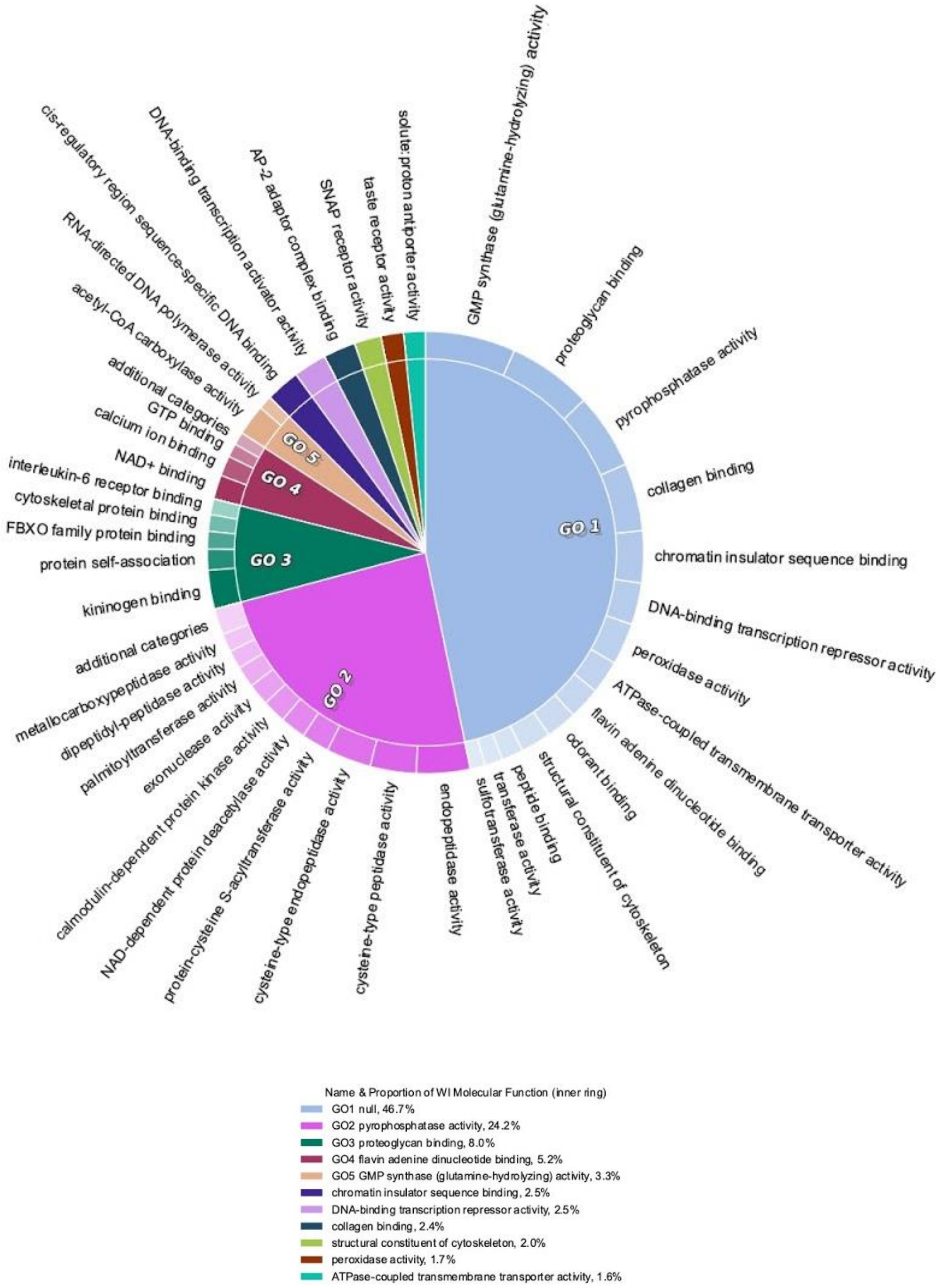


**Supplemental Figure S2**. Cirgo plots of significant GO terms by Molecular Function in HAN, WI.

**Evidence of hard-selective sweeps suggest independent adaptation to insecticides in Colorado potato beetle (Coleoptera: *Chrysomelidae*) populations**

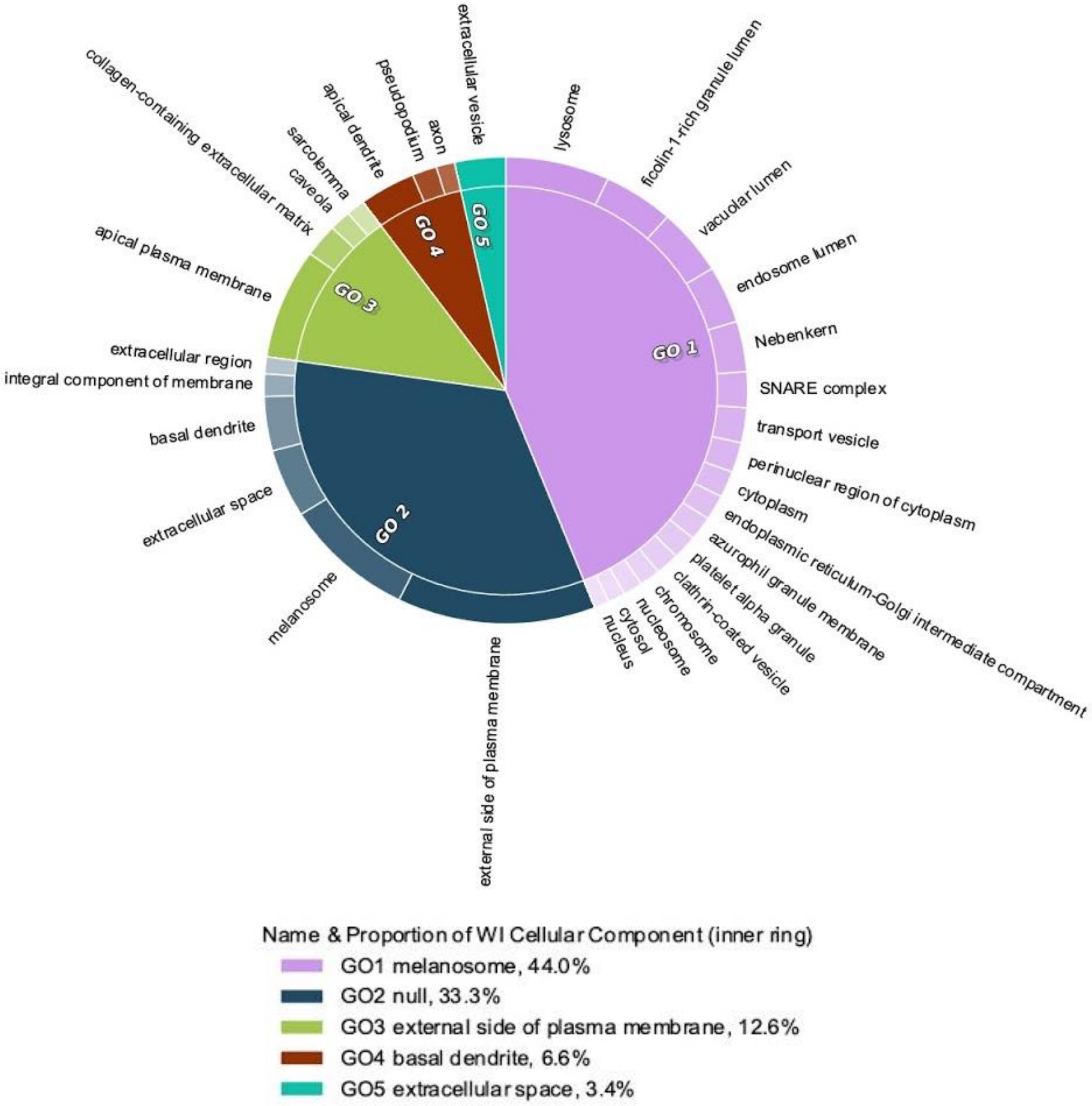


**Supplemental Figure S3**. Cirgo plots of significant GO terms by Cellular Component in HAN, WI.

**Evidence of hard-selective sweeps suggest independent adaptation to insecticides in Colorado potato beetle (Coleoptera: *Chrysomelidae*) populations**

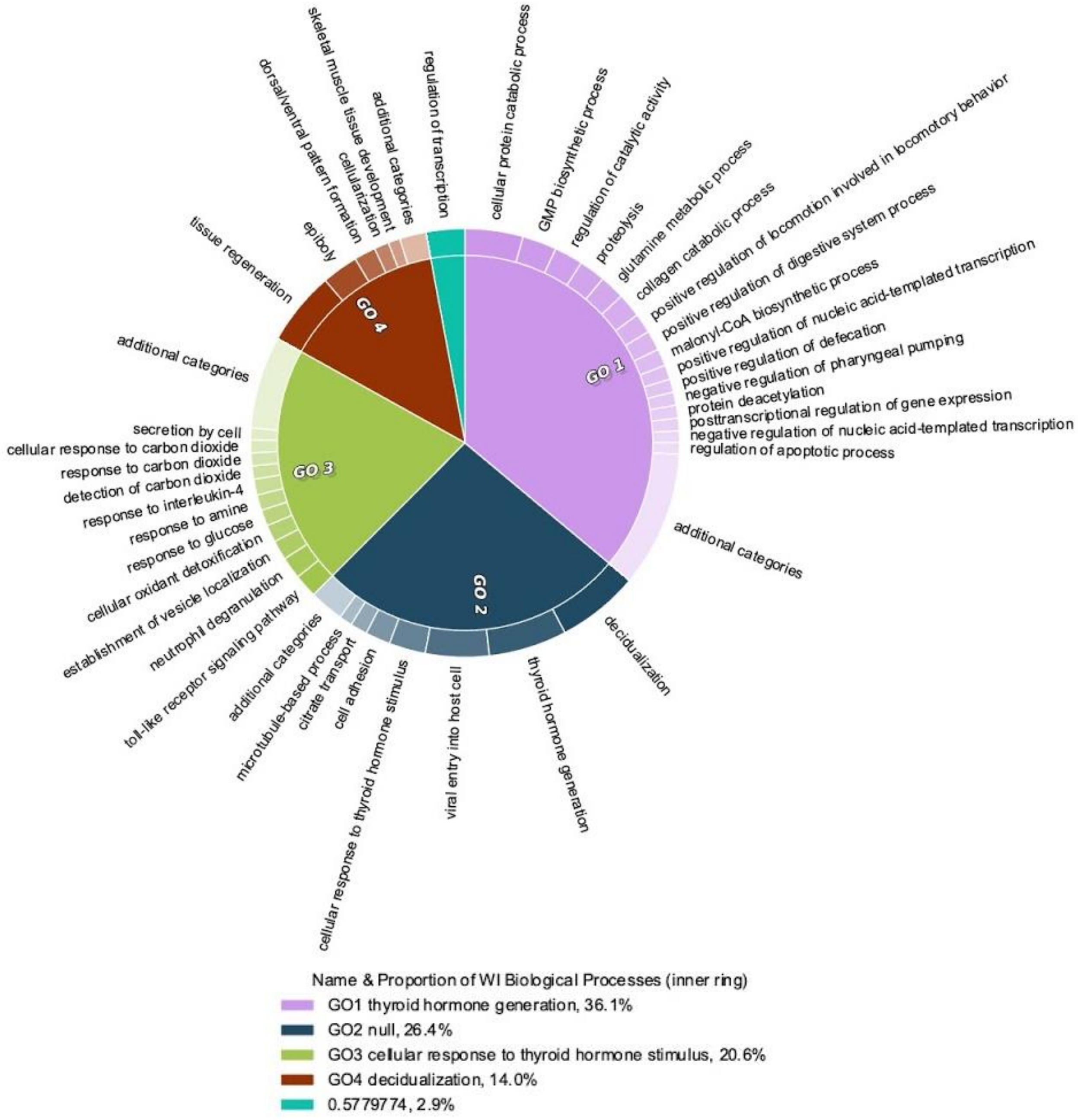


**Supplemental Figure S4**. Cirgo plots of significant GO terms by Biological Process in HAN, WI.

**Evidence of hard-selective sweeps suggest independent adaptation to insecticides in Colorado potato beetle (Coleoptera: *Chrysomelidae*) populations**

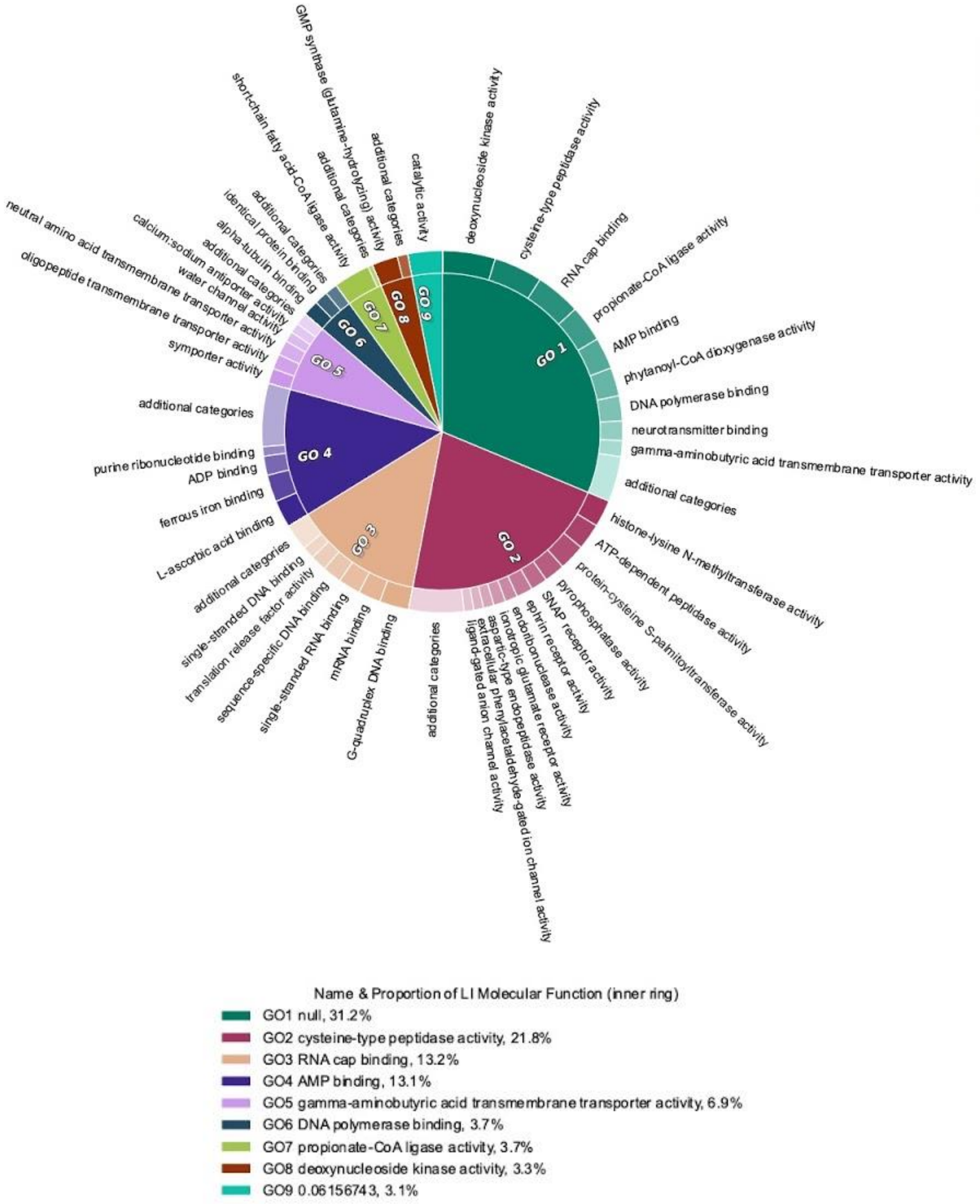


**Supplemental Figure S5**. Cirgo plots of significant GO terms by Molecular Function in Long Island, NY.

**Evidence of hard-selective sweeps suggest independent adaptation to insecticides in Colorado potato beetle (Coleoptera: *Chrysomelidae*) populations**

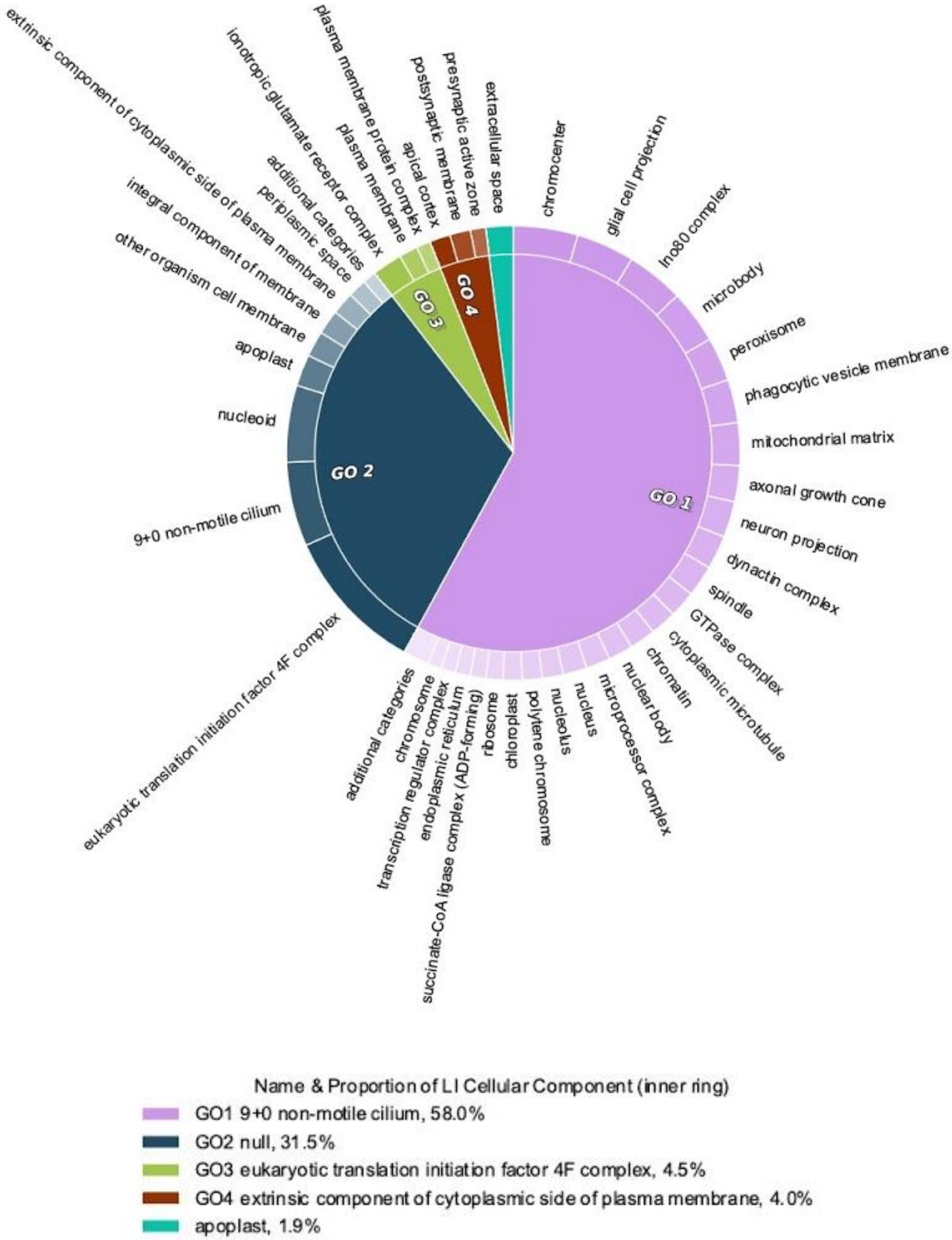


**Supplemental Figure S6**. Cirgo plots of significant GO terms by Cellular Component in Long Island, NY.

**Evidence of hard-selective sweeps suggest independent adaptation to insecticides in Colorado potato beetle (Coleoptera: *Chrysomelidae*) populations**

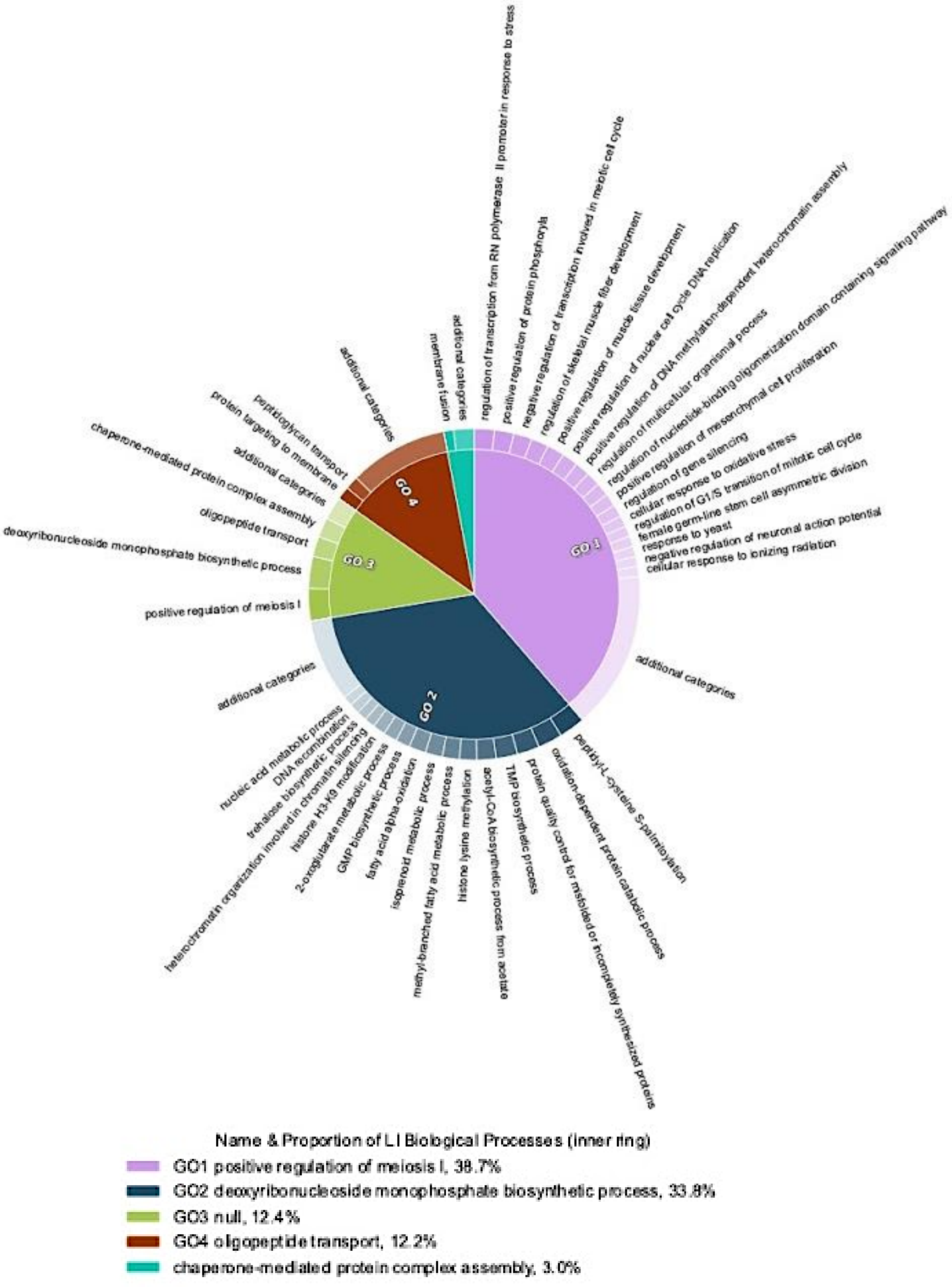


**Supplemental Figure S7**. Cirgo plots of significant GO terms by Biological Process in Long Island, NY.

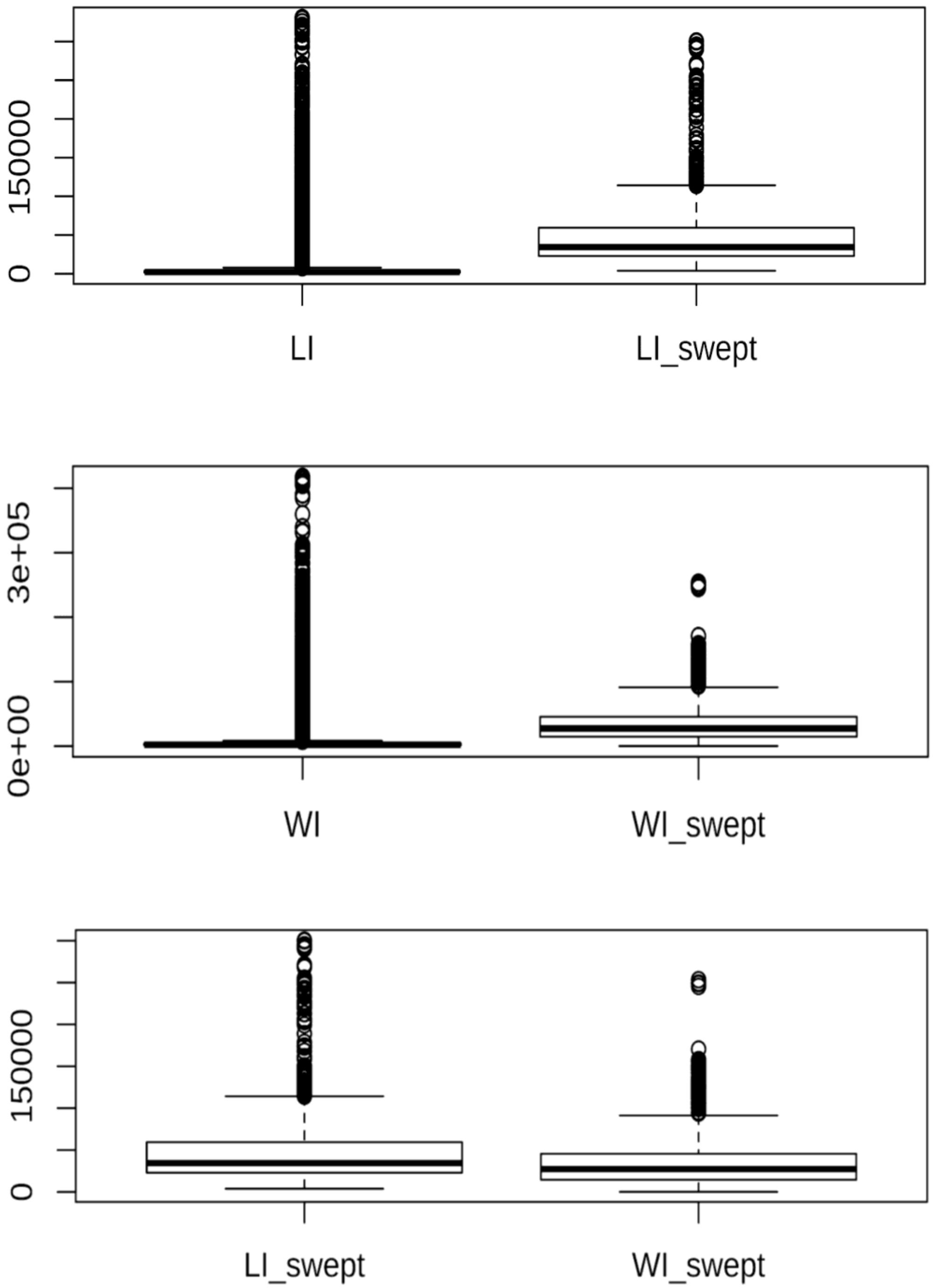


**Supplemental Figure S8.** Box and whisker plots for sweep region size (in base pairs) by population and type (swept and non-swept regions).

**Evidence of hard-selective sweeps suggest independent adaptation to insecticides in Colorado potato beetle (Coleoptera: *Chrysomelidae*) populations**
